# Clonally expanded CD38^hi^ cytotoxic CD8 T cells define the T cell infiltrate in checkpoint inhibitor-associated arthritis

**DOI:** 10.1101/2021.10.19.464961

**Authors:** Runci Wang, Anvita Singaraju, Kathryne E. Marks, Lorien Shakib, Garrett Dunlap, Amy Cunningham-Bussel, Lin Chen, Aidan Tirpack, Miriam R. Fein, Derrick J. Todd, Lindsey MacFarlane, Susan M. Goodman, Edward F. DiCarlo, Elena M. Massarotti, Jeffrey A. Sparks, Ole-Petter R. Hamnvik, Le Min, A. Helena Jonsson, Michael B. Brenner, Karmela K. Chan, Anne R. Bass, Laura T. Donlin, Deepak A. Rao

## Abstract

Immune checkpoint inhibitor (ICI) therapies that promote T cell activation have improved outcomes for advanced malignancies yet also elicit harmful autoimmune reactions. The T cell mechanisms mediating these iatrogenic autoimmune events remain unclear. Here we assayed T cells from joints of patients affected by ICI-induced inflammatory arthritis (ICI-arthritis), which can present clinically indistinguishable from rheumatoid arthritis (RA). Compared to the autoimmune arthritides RA and psoriatic arthritis (PsA), ICI-arthritis joints contained an expanded CD38^hi^ CD127^−^ CD8^+^ T cell subset that displays cytotoxic, effector, and interferon (IFN) response signatures. The abundance of CD38^hi^ CD8 T cells in ICI-arthritis resulted from a limited number of clones that could be found proliferating in the joint. Exposure of synovial T cells to Type I IFN, more so than IFN-γ, induces the CD38^hi^ cytotoxic phenotype. Relative to other CD8^+^ T cell subsets in the joints, the CD38^hi^ population is distinct from a dysfunctional population and clonally most related to TCF7^+^ memory populations. Examination of synovial tissue from bilateral knee arthroplasty demonstrated considerable sharing of TCR clonotypes in the CD38^hi^ CD8 T cell fraction from both knees. These results define a distinct CD8 T cell subset that may be directly activated by ICI therapy and mediate a tissue-specific autoimmune cellular reaction in patient joints.

## Introduction

Immune checkpoint inhibitors (ICI) augment T cell activation and have provided marked improvement in the outcome of multiple advanced cancers. Blocking immune checkpoint pathways such as PD-1/PD-L1 releases T cells from negative regulation and can induce potent anti-tumor responses ^(1)^. However, ICI therapy can also activate immune reactions against healthy tissues, leading to immune related adverse events (irAEs) in >80% of patients ^(1–8)^. These irAEs permit the study of human autoimmune responses from the unprecedented perspective of a defined inciting event—administration of ICI therapy. Defining the cellular events directly induced by ICI therapy and the downstream immune cascades may provide insights into mechanisms that maintain immune tolerance in the human immune system and prevent autoimmunity.

Here we focus on the inflammatory arthritis that develops following anti-PD-1/PD-L1 and anti-CTLA-4 therapies. This ICI-associated arthritis (ICI-arthritis) occurs in ~5% of treated patients and often clinically resembles classic inflammatory arthritides, such as rheumatoid arthritis (RA) and psoriatic arthritis (PsA), causing pain, swelling, and inflammatory joint effusions in both small and large joints ^(1, 3–7)^. ICI-arthritis can present within weeks-to-months after adminstering ICI therapy, but unlike most irAEs, can last years after therapy discontinuation. Joints affected create considerable pain, limit mobility and can become permanently damaged necessitating joint replacement surgery. ICI-arthritis is often treated with immunosuppressive therapies established for RA and PsA; thus, understanding the shared and distinct immune pathways in ICI-arthritis is critical for selecting effective therapies that target the adverse mechanisms. Further, an understanding of how the adverse mechanisms relate to the anti-tumor responses are needed to allow for opportunities to treat the adverse events and yet preserve cancer treatment.

The immune cell types active in the inflamed joints of ICI-arthritis are largely undefined, and the extent to which pathologic T cell responses are shared between ICI-arthritis and RA or PsA is unknown. Activated T cell infiltrates in RA and PsA joints identified through high dimensional analyses have emerged as defining features of these diseases. For seropositive RA joints, this includes the prominent expansion of CD4 T peripheral helper (Tph) cells, which provide local help to B cells and are marked by high levels of PD-1 ^(9)^. RA and PsA synovial fluid and tissue contain multiple subsets of activated CD4 and CD8 T cells ^(10–12)^, but lack large quantities of dysfunctional CD8 T cells such as found in the tumor microenvironment ^(13–17)^.

Here we directly compared T cell states found in the joints of patients with ICI-arthritis, RA, and PsA taking advantage of the ability to collect inflammatory joint fluid and isolate immune cells therein. Using mass cytometry and RNA-seq analyses, we identified a CD38^hi^ CD127^−^ CD8^+^ T cell population that is highly expanded in ICI-arthritis relative to the two classic idiopathic arthritides. These cells appear clonally expanded and exhibit features of cytotoxicity, proliferation, and activation. The T cell receptor (TCR) sequences are shared with a subset of cells that express *TCF7* and *IL7R*—implicating a possible stem-like and central memory progenitor for the ICI-induced autoimmune event. We further identified Type I interferons (IFN) as potential inducers of this T cell phenotype in ICI-arthritis.

## Results

### Mass cytometry identification of expanded CD38^hi^ CD8^+^ T cells in ICI-arthritis synovial fluid

To investigate the inflammatory features of ICI-arthritis, we analyzed mononuclear cells from synovial fluid of 6 ICI-arthritis, 5 seropositive RA and 5 PsA patients (**Supplementary table 1**) by mass cytometry using a panel that incorporated immunophenotyping markers for multiple cell types and specific T cell activation and effector states (**Supplementary table 2**). To measure PD-1 surface expression, we used a detection antibody that retains the capacity to bind PD-1 protein even when PD-1 is bound by a therapeutic anti-PD-1 antibody (**Fig. S1a**).

Within synovial fluid mononuclear cells, T cells were the largest population, accounting for ~50% of cells, followed by monocytes, NK cells, and then B cells. Comparable cell proportions were found across the 3 arthritides (**Fig. 1a, S1b**). The frequency of CD4 and CD8 T cells was also similar across the diseases (**Fig. 1a**). For the checkpoint pathway mediators PD-L1 and PD-L2, we found an increase in the portion of T cells and monocytes expressing these proteins in ICI-arthritis samples (**Fig. S1c**). In the RA joints, we observed an abundant PD-1^hi^ CXCR5^−^ CD4 T cell population, consistent with the Tph population previously described in RA (**Fig. S1d**) ^(9)^. The frequency of PD-1^hi^ Tph cells was substantially lower in ICI-arthritis, comparable to the frequencies seen in PsA joints (**Fig. S1d**).

**Figure 1.**
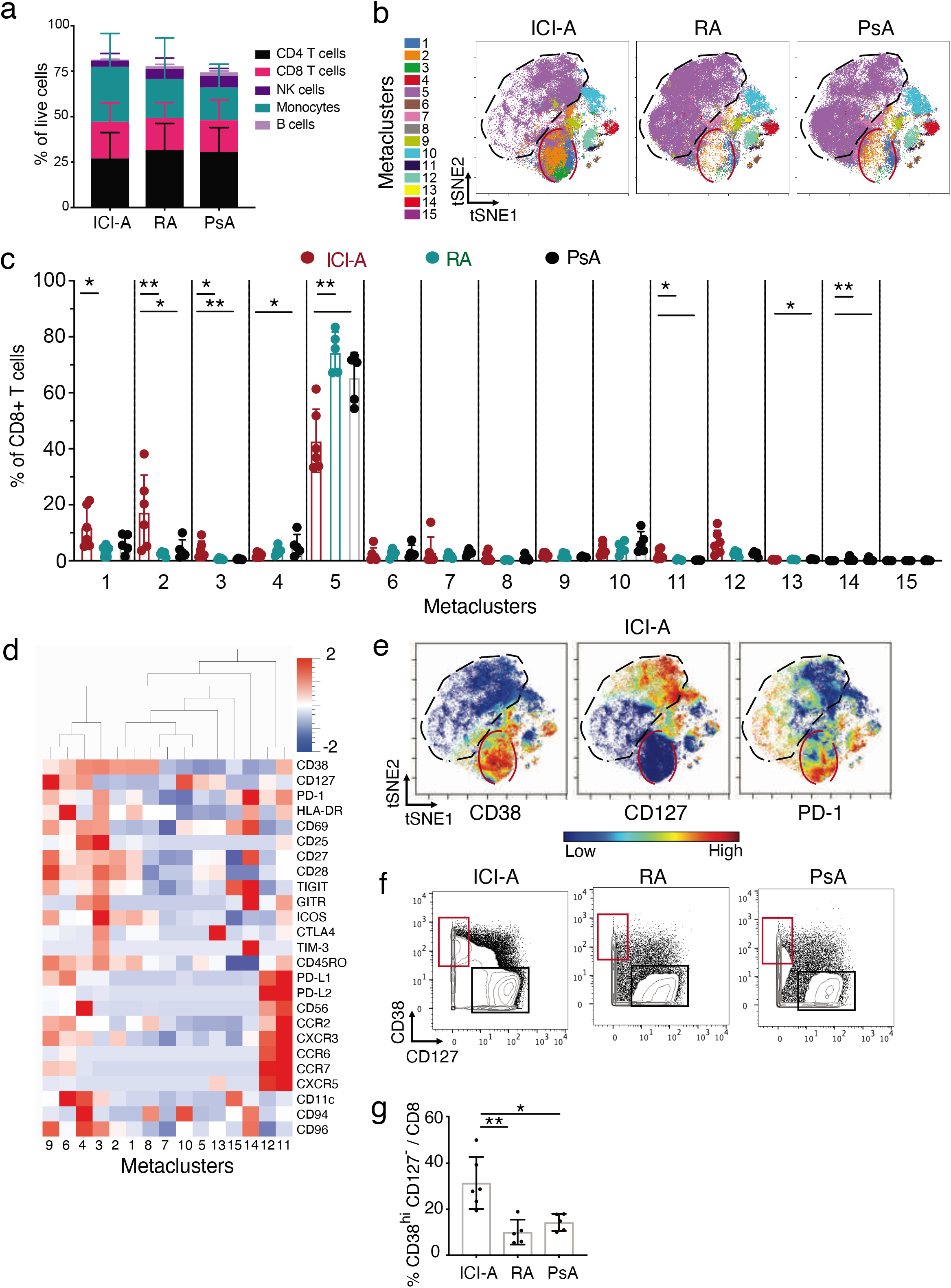
Expansion of CD38^hi^CD127^−^ CD8 T cells in ICI-arthritis (ICI-A). **(a)** Frequency of CD8 T cells, CD4 T cell, B cells, monocytes, NK cells and T cells in ICI-A (n=6), RA (n=5) and PsA (n=5) synovial fluid. **b)** tSNE visualization of FlowSOM metaclusters of CD8 T cells from ICI-A, RA and PsA synovial fluid. The red circle indicates area that includes MC1-3 with increased density in ICI-A. Black circle indicates area of MC5 with lower density in ICI-A. **c)** Frequency of FlowSOM metaclusters of CD8 T cells from ICI-A, RA and PsA synovial fluid. d) Heatmap of marker expression in CD8 T cell metaclusters. **e)** tSNE plots of mass cytometry data showing expression of indicated markers on CD8 T cells from ICI-A, RA and PsA synovial fluid. Red and black circles are shown as in (b). **f,g)** Biaxial gating (f) and quantification (g) of CD38^hi^ CD127^−^ cells among CD8 T cells from ICI-A, RA and PsA synovial fluid detected by mass cytometry. Mean ± SD shown. * p<0.05, **p<0.001, *** p<0.0001 by Kruskal-Wallis test in (c) and (g).

To further explore differences in CD8 T cell phenotypes, we used FlowSOM to define metaclusters (*i.e*. populations) among synovial fluid CD8 T cells and compare their metacluster frequencies across arthritides (**Fig. S2a**). This analysis revealed a significant expansion of metaclusters 1-3 (MC 1-3) in ICI-arthritis samples compared to RA (5-fold increase) and PsA (3-fold increase) samples (**Fig. 1b,c**). Analysis of marker expression revealed a shared CD38^hi^ CD127^−^ expression pattern across MC1-3, which together comprised ~33% of CD8 T cells in ICI-arthritis samples (**Fig. 1d-f**). Biaxial gating on CD38^hi^ CD127^−^ cells validated the unbiased clustering result to confirm significant expansion of CD38^hi^ CD127^−^ cells in ICI-arthritis (**Fig. 1g**). Conversely, MC5, containing CD38^−^ cells, was substantially reduced in ICI-arthritis (42%) compared to RA (75%) and PsA (65%) (**Fig. 1c-e**). Among the smaller clusters, MC11 (2%, CD38^+^ CD127^−^ PD-1^+^ PD-L1^+^ PD-L2^+^) was increased in ICI-arthritis, while MC4 (2%, CD38^+^ CD127^+^ PD-1^+^ CD56^+^), MC13 (0.3%, CTLA-4^+^ CXCR5^+^), and MC14 (0.01%, PD-1^hi^ TIM-3^+^ TIGIT^+^) were decreased (**Fig. 1c,d**).

Cells in MC 1-3 shared expression of HLA-DR, CD45RO, CD27, CD28, TIGIT and ICOS (**Fig. 1d**). Among these 3 MCs, MC3 showed the highest expression of activation-associated markers, including CD69, CD25, PD-1, TIGIT, GITR, ICOS, CD96, CTLA-4 and TIM-3, whereas MC1 and MC2 were negative for GITR, CTLA-4 and TIM-3 and expressed lower levels of CD69, CD25, and PD-1 (**Fig. 1d, S2b**). These results suggest that a collection of CD38^hi^ CD127^−^ CD8 T cells with a range of activation marker expression is selectively expanded in ICI-arthritis.

### CD38^hi^ CD127-CD8 T cells in ICI-arthritis express cytotoxic effector and proliferative programs

To evaluate the transcriptional programs that characterize T cells from ICI-arthritis synovial fluid, we sorted 5 CD8 T cell populations from 7 ICI-arthritis joint samples for bulk RNA-seq, as well as the same T cell populations from 6 seropositive RA and 6 PsA samples for comparison (**Fig. 2a, S3a,b, Supplementary table 1**). The CD8 populations included CD38^hi^ CD127^−^ PD-1^int^ cells, as well as comparator populations including PD-1^int^ or PD-1^−^ CD38^−^ CD127^+^ cells (to capture the largest comparator population), PD-1^hi^ cells (to capture dysfunctional cells), and CD38^−^ CD127^−^ KLRG1^+^ cells (to capture cytotoxic cells). Consistent with mass cytometry results, quantification of this flow cytometry data demonstrated that CD38^hi^ CD127^−^ cells were the largest population in ICI-arthritis joints, followed by CD38^−^ CD127^+^ PD-1^int^ cells and CD38^−^ CD127^+^ PD-1^−^ cells (**Fig. 2b**).

**Figure 2.**
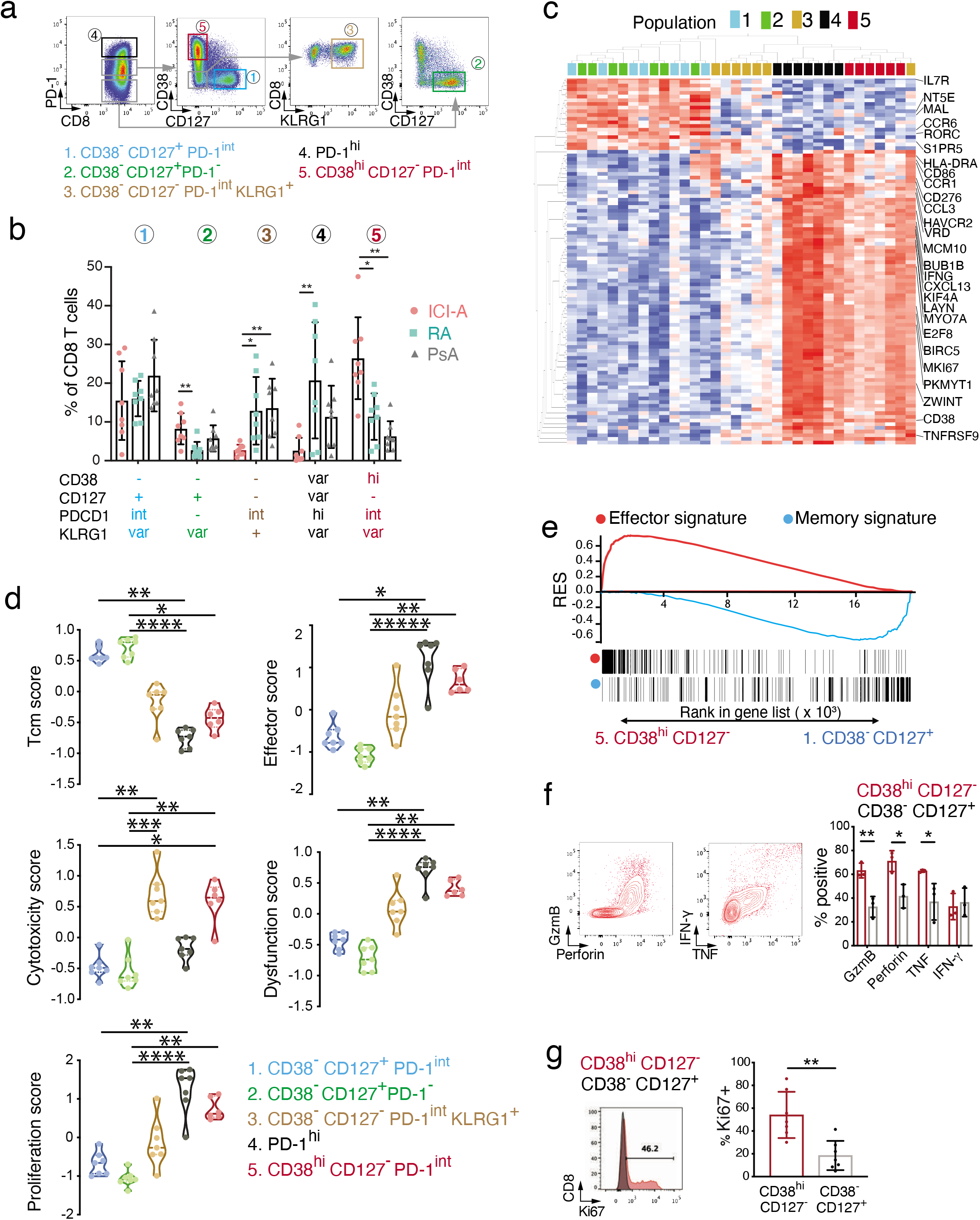
Transcriptomic features of CD8 T cells in ICI-arthritis synovial fluid. **a)** Sorting scheme to isolate 5 CD8 T cell populations of interest. **b)** Frequency of sorted CD8 T cell populations in ICI-arthritis (ICI-A), RA and PsA synovial fluid (n=8). **c)** Hierarchical clustering of CD8 T cell populations from CI-A synovial fluid based on row-normalized mean expression of 100 top variable genes by RNA-seq. Colors indicate cell populations as in (a). **d)** Gene module scores of CD8 T cell populations as in a) from CI-A synovial fluid, calculated based on differentially expressed genes. **e)** Distribution of genes in the effector signature gene set (red line, GOLDRATH EFF_VS_MEM_CD8_TCELL_UP) and memory signature gene set (blue line, GSE9650_EFFECTOR_VS_MEMORY_CD8_TCELL_DN) in sorted CD38^hi^ CD127^−^ and CD38^−^ CD127^+^ populations from CI-A synovial fluid, plotted with running enrichment score (RES) and ranks in gene list detected by RNA-seq. **f)** Representative flow cytometric plots and summarized frequency of intracellular granzyme B, perforin, TFN-y, and TNF in CD38^hi^ CD127^−^ and CD38^−^ CD127^+^ populations sorted from ICI-A synovial fluid, detected after PMA/ionomycin stimulation (n=3 donors). **g)** Representative flow cytometric plot and frequency of intracellular Ki67 in sorted CD38^hi^ CD127^−^ and CD38^−^ CD127^+^ populations from ICI-A synovial fluid (n=7). Mean ± SD shown. * p<0.05, **p<0.001, *** p<0.0001 by Kruskal-Wallis test in (b), (d), (f) and (g).

From the ICI-arthritis samples, transcriptomic comparison across the 5 CD8 T cell populations identified 1,809 genes differentially expressed (FDR q value<0.01, **Supplementary table 3**). Evaluation of the top 100 most variable genes revealed distinct expression patterns, clearly separating the CD38^hi^ CD127^−^ cells and PD-1^hi^ cells from the CD38^−^ CD127^+^ cells (**Fig. 2c**). Upregulated in the CD38^hi^ CD127^−^ cells and PD-1^hi^ cells were genes expressed upon activation (*IFNG*, *HLA-DRA*, *CD86*, *CD38*, *TNFRSF9*), dysfunction (*HAVCR2*, *CXCL13*), and proliferation (*MCM10*, *BUB1B*, *KIF4A*, *MYO7A*, *BIRC5*, *MKI67*, *PKMYT1, ZWINT)*. In contrast, CD38^−^ CD127^+^ cells preferentially expressed genes associated with memory states (*IL7R*, *MAL, CCR6*). KLRG1^+^ cells showed no strong enrichment for these memory associated genes.

To further investigate these differences, we compiled gene lists summarizing modules of memory and effector states, cytotoxicity, dysfunction, and proliferation from published studies and calculated scores based on the average expression for each cell population (**Supplementary table 4**) ^(14–21)^. These 5 modules were examined by 2 approaches. First, the full list was used for scoring to evaluate broad expression patterns (module score, **Fig. S4a**). Second, the differentially expressed genes identified by ANOVA were assigned to these modules and used for scoring to evaluate the difference in expression among populations (DEG score, **Fig. 2d**). In both cases, the CD38^hi^ CD127^−^ cells showed high expression of effector, cytotoxicity, dysfunction and proliferation modules and low expression of the memory module. PD-1^hi^ cells showed a similarly high expression of effector, dysfunction and proliferation modules but low cytotoxicity relative to CD38^hi^ CD127^−^ cells. In contrast, CD38^−^ CD127^+^ populations showed a transcriptomic pattern characteristic of central memory T cells, as opposed to effector memory T cells (Tem), CD45RA^+^ effector memory T cells (Temra) or resident memory T cells (Trm) (**Fig. S4a**), suggesting that even cells with intermediate PD-1 expression can appear primarily as central memory cells by global transcriptomic analysis. CD38^−^ CD127^+^ cells also showed low effector, cytotoxicity, dysfunction, and proliferation scores compared to CD38^hi^ CD127^−^ cells and KLRG1^+^ cells, suggesting that downregulation of CD127 may help distinguish memory from effector and dysfunctional states independently of CD38 level. Despite very different PD-1 expression, the CD38^−^ CD127^+^ PD-1^int/-^ cells showed lower cytotoxicity features similar to that of PD-1^hi^ cells, setting these 3 populations apart from the more cytotoxic appearing CD38^hi^ CD127^−^ and KLRG1^+^ cells (**Fig. 2d, S4a**).

We also conducted pairwise comparisons between each CD8 T cell population. These analyses similarly demonstrated higher expression of genes associated with a Tcm phenotype, including *IL7R, TCF7, CCR7,* and *SELL,* in CD38^−^ CD127^+^ PD-1^−^ cells and CD38^−^ CD127^+^ PD-1^int^ cells, higher expression of proliferation associated genes (*AURKB, CCNB2, CENPN, KIF2C/15/18B, MCM5/10, ZWINT*) in CD38^hi^ CD127^−^ and PD-1^hi^ populations, and higher expression of cytotoxicity associated genes (*GZMB*, *GZMH*, *FGFBP2)* in CD38^hi^ CD127^−^ and KLRG1^+^ cells (**S4b**). These populations also exhibited differential expression of chemokines and chemokine receptors, indicating potential differences in migratory capacity (**Fig. S4c**).

To further examine the difference in the two largest populations: the expanded CD38^hi^ CD127^−^ PD-1^int^ cells versus CD38^−^ CD127^+^ PD-1^int^ cells, gene set enrichment analysis was performed to identify enriched MSigDB pathways. This analysis identified 250 pathways enriched in CD38^hi^ CD127^−^ PD-1^int^ cells and 21 pathways enriched in CD38^−^ CD127^+^ PD-1^int^ cells (FDR <0.05). Among these, multiple gene sets associated with proliferation were enriched in CD38^hi^ CD127^−^ PD-1^int^ cells (**Supplementary table 5**). In addition, multiple pathways associated with active T cell effector functions were enriched in CD38^hi^ CD127^−^ cells, while pathways associated with naïve or quiescent memory states were among those enriched in CD38^−^ CD127^+^ cells. As one example, opposing signatures for effector (Goldrath Effector vs memory CD8 T cell UP) and memory (GSE9650 effector vs memory CD8 T cell DN) were enriched in CD38^hi^ CD127^−^ cells and CD38^−^ CD127^+^ cells respectively (**Fig. 2e**), confirming the findings identified by differentially expressed genes.

To evaluate protein expression levels for cytotoxic mediators, cells subsets were sorted and stained for intracellular granzyme B, perforin, IFN-*γ*, and TNF after stimulation with PMA/ionomycin. Compared to CD38^−^ CD127^+^ cells, CD38^hi^ CD127^−^ cells more frequently expressed granzyme B, perforin, and TNF, consistent with RNA-seq results (**Fig. 2f**). Further, CD38^hi^ CD127^−^ cells more frequently expressed Ki67, consistent with increased proliferation (**Fig. 2g**).

### CD38^hi^ CD8 T cells are clonally expanded and actively proliferating in ICI-arthritis joints

To evaluate CD8 T cells within ICI-arthritis joints with higher resolution and in an unbiased manner, we performed single-cell transcriptomic (scRNA-seq) and antigen receptor sequencing on sorted CD8 T cells from synovial fluid from four ICI-arthritis patients and from synovial tissue from both knees of an additional ICI-arthritis patient (**Fig. S5a**). From 18,472 CD8 T cells that passed quality filtering (<3,000 total genes, <10% mitochondrial genes), we defined 8 transcriptionally distinct clusters (**Fig. 3a, S5b,c**). Cluster 0, the most abundant cluster, featured high expression of *HLA-DRA*, *CD74*, *IFNG*, and multiple granzymes including *GZMA*, *GZMB*, *GZMH* and *GZMK*. In contrast, Clusters 1, 2, and 4 showed higher expression of central memory markers including *IL7R* (CD127), *TCF7*, *SELL*, and *CCR7*. Cluster 4 was further distinguished by expression of ZNF683, a tissue-resident memory marker, and a strong IFN signature. Cluster 3 contained cells with *KLRB1*, *RORC*, *ZBTB16*, and TCR chains *TRAV1-2* and *TRBV6-4,* consistent with a mucosal-associated invariant T (MAIT) cell phenotype. Cluster 5 contained a mixture of gamma/delta T cells and CD8 T cells expressing a combination of effector markers such as *KLRG1*, *KLRC1/2, KLRD1* and *XCL1/2*. Cluster 6 was distinguished by high expression of genes associated with dysfunction, including *PCDC1, ENTPD1, CTLA4, HAVCR2,* and *TIGIT.* Cluster 7 captured proliferating cells, marked by strong expression of cell-cycle genes including *MKI67, ZWINT, CENPF,* and *PCNA* (**Fig. 3b**).

**Figure 3.**
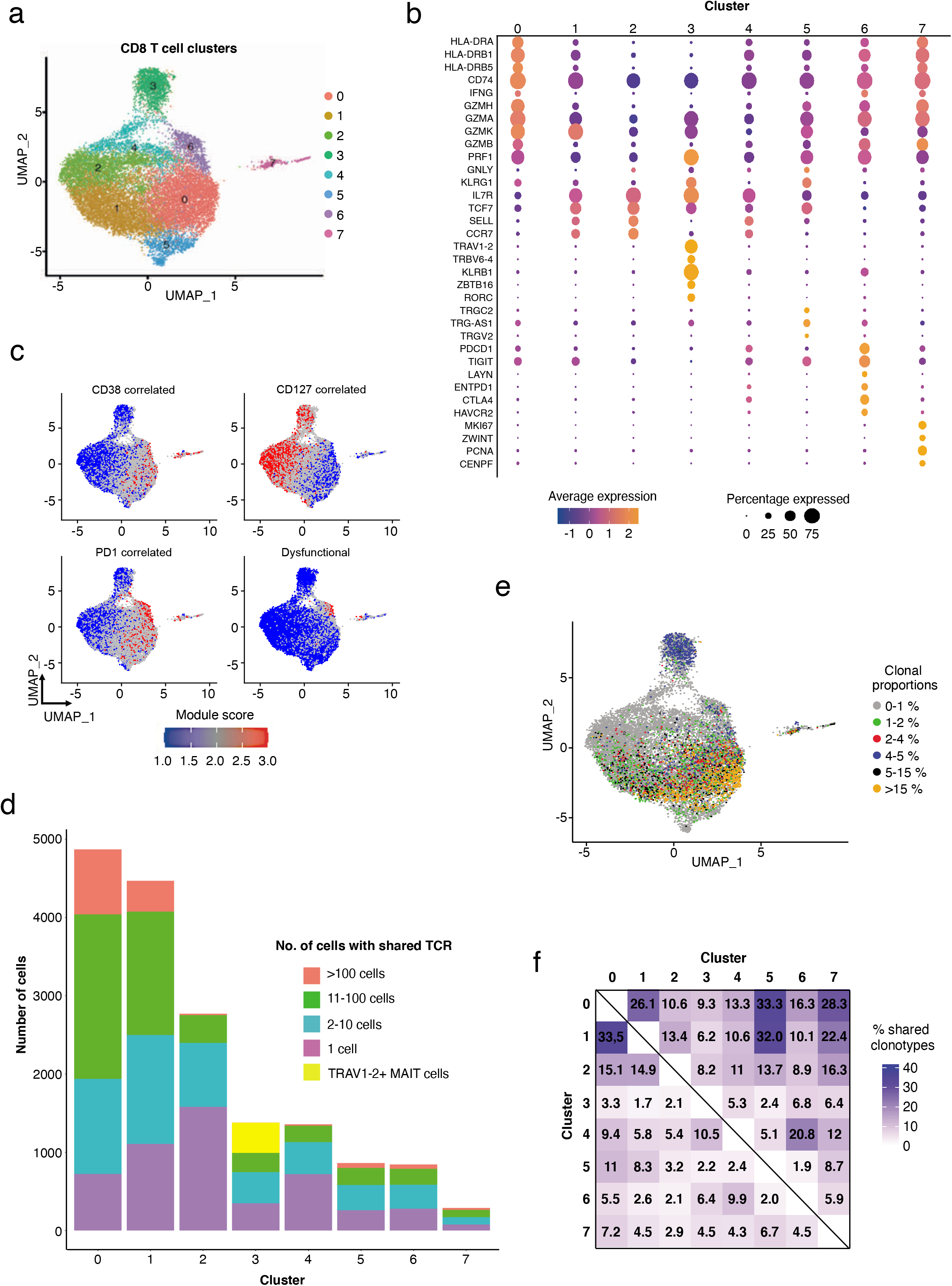
Extensive clonal expansion of synovial CD38^hi^ CD127^−^ CD8 T cells in ICI-arthritis. **a)** Synovial fluid (n=4) and tissue (bilateral knee explants from n=1) CD8 T cells analyzed by scRNAseq and plotted in clusters in UMAP space. **b)** Dotplot visualization of differentially expressed genes that distinguish CD8 T cell clusters. Color of the dot represents the average expression of the gene across cells in the cluster. Size of the dot represents the percentage of cells in the cluster with that gene detected. **c)** UMAP overlay of signature scores for gene sets that correlate with CD38, IL7R (CD127) or PDCD1 (PD-1) expression derived from bulk RNA-seq data, and a CD8 T cell dysfunctional module. **d)** The number of cells in each cluster that either contains a unique TCR (purple) or a shared TCR (all other colors, for which the total number of clones from the sample is represented by the specific color). Yellow in Cluster 3 represents TRAV1-2+ invariant MAIT cells. **e)** Highly expanded TCR clones (>1%) from each patient depicted onto the transcriptionally-defined UMAP clustering. **f)** The percentage of clonotypes shared between clusters. The row identity represents the denominator.

In Cluster 0, the high level of expression of *CD74, HLA-DRA, GZMB, GZMH*, and *GZMK*, and the distinct absence of *IL7R* raised the possibility that these cells represented the CD38^hi^ CD127^−^ population initially identified by mass cytometry. The expression of the CD38 gene itself was poorly detected in the scRNA-seq data, yet CD38 protein was detected during flow sorting on ~30% of CD8 T cells in these samples (**Fig. S5a**). To identify potential CD38^hi^ CD127^−^ cells in the single-cell transcriptomic dataset, we adopted the following approach. Using our bulk RNA-seq data, we first generated two gene modules: (i) genes with expression patterns that correlated with *CD38* expression across all CD8 populations (Pearson correlation coefficient r > 0.6) (**Supplementary table 6**) and (ii) genes with an expression pattern in CD38^hi^ CD127^−^ cells that distinguished these cells from the other CD8 T cell populations analyzed by bulk RNA-seq (**Supplementary table 7**). We then calculated the module scores for each CD8 T cell in the scRNA-seq data. In both instances, cells in Cluster 0 showed the highest score for CD38^hi^ cells, as visualized on the UMAP (**Fig. 3c, S5d**). In contrast, Clusters 1 and 2 displayed the opposite pattern, with low scores for CD38^hi^ CD127^−^ module and high scores for a CD38^−^ CD127^+^ module, and high levels of *IL7R* expression (**Fig. 3c, S5d, Supplementary table 7**). These analyses support that the cells in Cluster 0 represent the CD38^hi^ CD127^−^ PD-1^int^ cells identified by mass cytometry and bulk-sorted RNA-seq. Thus, across 3 distinct cellular analysis pipelines and 17 ICI-arthritis patients, we consistently identified an enrichment of the CD38^hi^ CD127^−^ population in ICI-arthritis joints.

In the scRNA-seq analysis, Cluster 6 displayed the highest expression of genes associated with the PD-1^hi^ sorted bulk T cell transcriptome (**Fig. S5d, Supplementary table 7**). Cluster 6 cells also scored highest for a module of genes correlated with *PDCD1* expression, which was notably lower in Cluster 0 cells (**Fig. 3c, Supplementary table 6**). Additionally, we probed the expression of a CD8 T cell dysfunctional module derived in the context of a tumor microenvironment (22) and found that Cluster 6 cells scored highest for this module as well (**Fig. 3c, Supplementary table 7**). Taken together, gene expression analysis at the single-cell resolution supports a clear distinction between the CD38^hi^ CD127^−^ cells in Cluster 0 and dysfunctional cells in Cluster 6, despite similar expression patterns for effector cell factors such as *GZMB, GZMH, GZMK, GZMA* and *IFNG* (**Fig. 3b**). Relative to Cluster 0, the Cluster 6 dysfunctional cells represent a relatively small portion (4.8 %) of CD8 T cells in ICI-arthritis synovial fluid.

We then used the T cell receptor (TCR) sequences from each CD8 T cell to examine clonal distributions within and across the transcriptionally-defined clusters (**Fig. 3d**). The Cluster 2 memory population contained the highest number and frequency of cells with a unique TCR sequence (~1,500, 60%). In contrast, despite having almost twice as many total cells, Cluster 0 CD38^hi^ contained fewer T cells with a unique TCR (~900, 20%) and instead exhibited extensive clonal expansion of a smaller number of TCR clonotypes (~600 clonotypes found across ~4000 cells). The six largest clonotypes in Cluster 0 each contained over 100 T cell clones. Notably, the frequency of shared clones in Cluster 0 surpassed even the semi-invariant TCR MAIT population of Cluster 3. From the perspective of individual patients, the most expanded clonotypes for each patient, that is, those accounting for >5% of all T cells, were concentrated primarily in Cluster 0, with some representation in Cluster 1 as well (**Fig. 3e**). These observations are consistent with Clusters 0 and 3 exhibiting the lowest score for repertoire diversity as measured by the Simpson Index and Shannon Equitability metric, which account for variability in abundance and evenness (**Fig. S5e**). Together these data indicate the large number of T cells with a CD38^hi^ signature resulted from extensive clonal expansion.

We next sought to understand relationships between transcriptionally-defined clusters from the TCR clonotype perspective. Cluster 0 CD38^hi^ cells were most highly related to Cluster 1 Tcm-like, Cluster 5, and Cluster 7 proliferating cells; with ~30% of Cluster 0 TCR clonotypes shared in each of these clusters (**Fig. 3f**). The dysfunctional CD8 T cells in Cluster 6 shared the highest proportion of TCR clonotypes with the memory Cluster 4 population (10%). The actively proliferating Cluster 7 cells had the highest frequency of TCR clonotypes in Cluster 0 and Cluster 5 (7.2% and 6.7%, respectively). Cluster 2, a memory population, had a large number of unique clones, yet had the highest frequency of TCR clone overlap with Cluster 0, 1, 5 and 7. Cluster 2 expressed a central memory signature and had the highest expression of *TCF7*, suggesting the potential for a stem cell memory population. As Cluster 2 and 1 are largely transcriptionally similar, with notable differences in Cluster 1 including higher levels of functional effector features such as *GZMK*, this may suggest a transitional state between the stem cell memory-like Cluster 2 and the activated Clusters 0 and 5.

In summary, scRNA-seq of synovial CD8 T cells confirmed the presence of the unique, expanded CD38^hi^ CD127^−^ effector population, both in the fluid and tissue in ICI-arthritis, which appears oligoclonal and distinct from dysfunctional T cells.

### Considerable overlap of TCR clones across right and left knee in ICI-arthritis

A patient requiring bilateral total knee replacements after ICI therapy provided the opportunity to probe CD8 T cells in the synovium across both knees of the same individual. The aggressive potential of ICI-arthritis was particularly notable in this patient, who had no underlying autoimmune condition nor autoantibodies consistent with RA. Within 6 months of ICI therapy, which cleared their metastatic melanoma, the patient experienced joint swelling and pain and, within 2 years, they required replacement of both knees. Radiographs of the knees showed symmetrical joint space narrowing of both medial and lateral compartments and periarticular osteopenia, consistent with an inflammatory arthritis (**Fig. 4a**). Histologically, the synovium from the left knee was scored as an acute and chronic inflammatory reaction, involving mostly mononuclear cell and neutrophil infiltrates. The right knee, operated on 6 months after the left, was histologically indistinguishable from RA with expansive lymphocyte aggregates and plasma cell communities, Russel bodies and binucleate plasma cells (**Fig. 4b,c**) ^(23)^. At the single-cell and molecular level, the synovial tissue CD8 T cells exhibited transcriptional phenotypes and ratios comparable to that of ICI-arthritis synovial fluid, including a predominance of the CD38^hi^ CD127^−^ effector population (**Fig. 4d, S5a**). Strikingly, TCR-sequencing demonstrated considerable sharing of TCR clonotypes between the left and right knees, with a predominance of shared clonotypes in Cluster 0 (16%) and Cluster 5 (12%), which contrasted minimal sharing of memory cell clonotypes from Cluster 2 and 4 (3 and 1%, respectively) (**Fig. 4e**).

**Figure 4.**
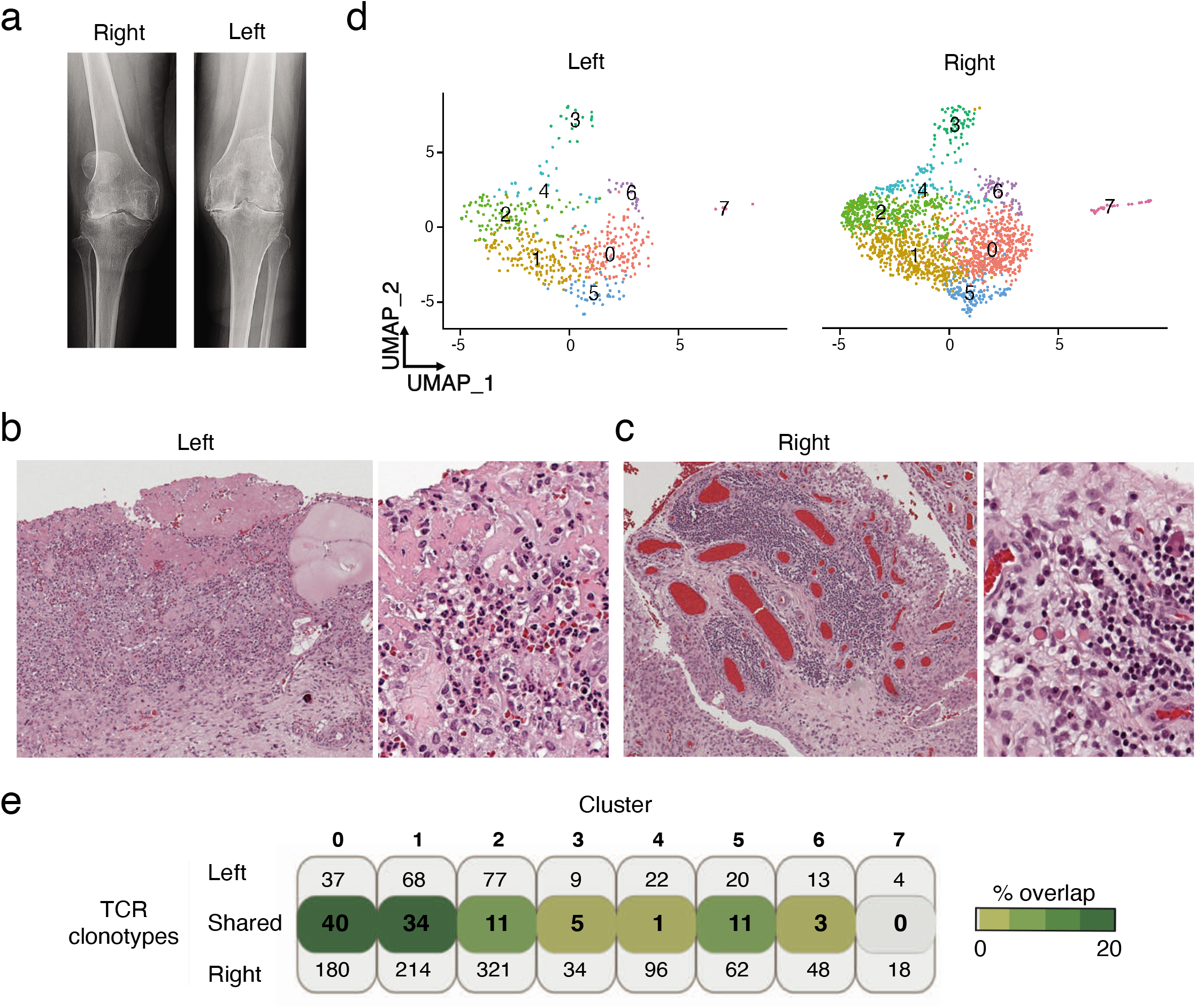
Bilateral knee ICI-arthritis with extensive inflammation and matching TCR clonotypes. **a)** Left and right knee radiographs in preparation for joint replacement surgery. **b)** Left knee synovial tissue H&E histology with extensive neutrophil and mononuclear infiltration, as well as fibrin exudate into synovial cavity. **c)** Right knee synovial tissue with extensive lymphocyte aggregate formation with plasma cell communities and Russel bodies. **d)** Synovial tissue CD8 T cell scRNA-seq clusters from the left and right knee. **e)** Number of distinct CD8 TCR clonotypes in left and right knee from each transcriptionally-defined cluster. The percent of overlap of the total cells from each cluster is depicted in shades of green with the darkest color indicating the highest frequency of overlap.

### Distinct activation of IFN-inducible genes in ICI-arthritis

To identify transcriptomic features that distinguish T cells from ICI-arthritis samples as compared to RA and PsA more broadly, we computationally combined bulk RNA-seq data from all sorted populations from each disease and tested for altered pathways comparing ICI-arthritis to RA+PsA. Nine reactome pathways were enriched and four pathways were decreased in ICI-arthritis CD8 T cells as compared to CD8 T cells from RA and PsA (**Supplementary table 8**). Among the pathways upregulated in ICI-arthritis samples were both Type I and Type II IFN pathways. Consistent with this result, many IFN-inducible genes were detected among the list of genes differentially expressed between ICI-arthritis T cells and combined RA+PsA T cells (**Supplementary table 9**). When examined in pair-wise comparisons of ICI-arthritis vs RA and ICI-arthritis vs PsA, IFN-inducible genes such as *OAS1/3, STAT1/2, IFIT1/3, CXCL9/10/11, IFI35/44/44L, IRF1/7/9, ISG15,* and *MX1* were significantly upregulated in ICI-arthritis T cells (**Fig. S6a**). To investigate the extent of IFN signaling, we calculated a score reflecting expression of a set of 106 IFN-inducible genes (**Supplementary table 4)**. ICI-arthritis samples showed marked upregulation of this gene set, indicating a pattern of broad IFN activation in ICI-arthritis (**Fig. 5a**).

**Figure 5.**
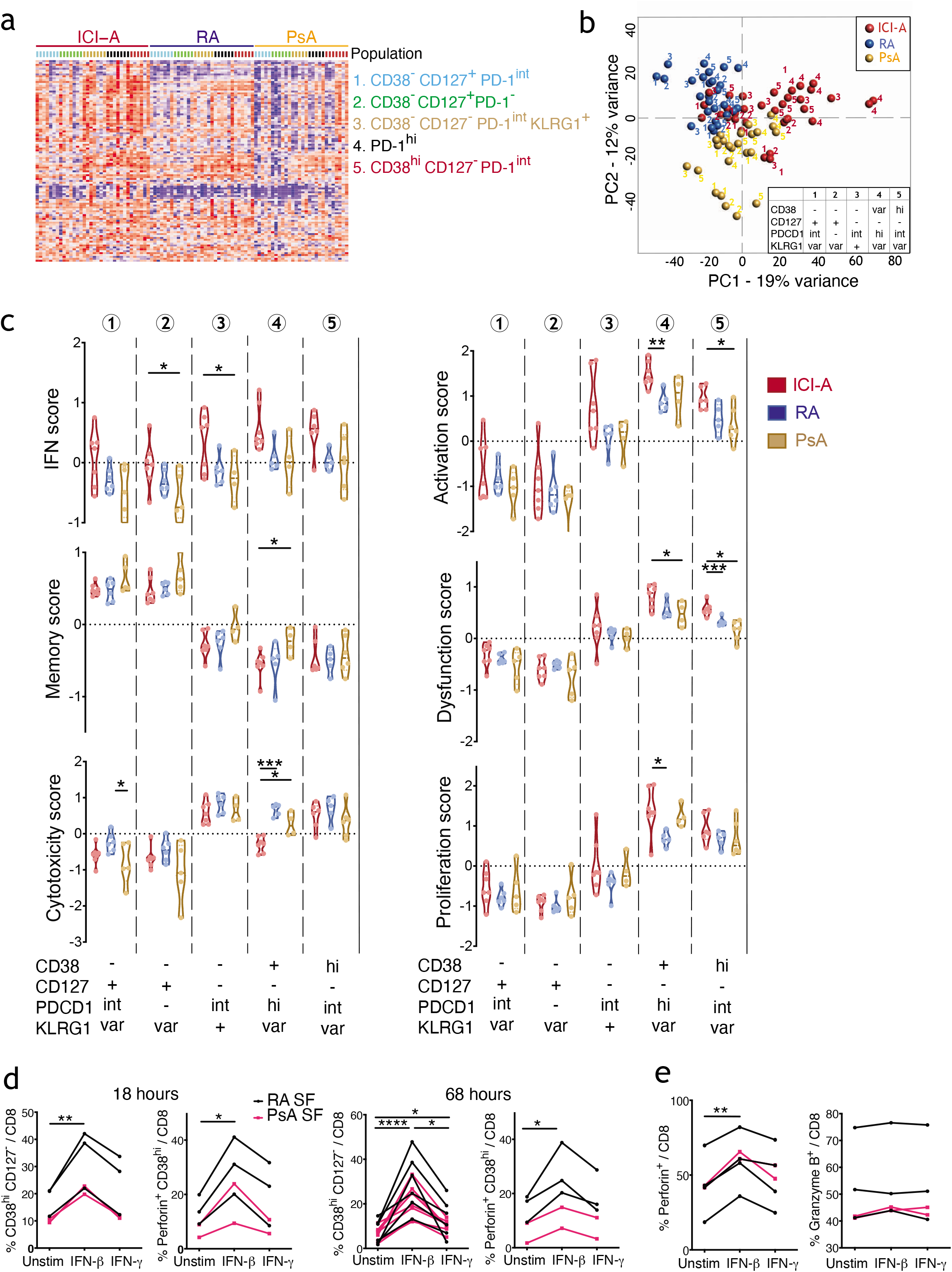
A Type I IFN signature in CD8 T cells in ICI-arthritis. **a)** Heatmap showing the expression of 106 IFN-inducible genes across the sorted populations ordered by diseases. b) PCA plot showing clusters of CD8 T cell populations using genes differentially expressed by ICI-A, RA and PsA, with numbers indicating 5 sorted populations colored by disease. **c)** Gene module scores of the sorted CD8 T cell populations as in c-d) from ICI-A, PsA and RA synovial fluid, calculated based on differentially expressed genes. **d,e)** Frequency of CD38^hi^ CD127^−^ cells and CD38^hi^ perforin+ cells (d), and perforin^+^ cells and granzyme B+ cells (e) in CD8 T cells from RA or PsA SFMC cultured with IFN-β or IFN-γ for indicated times. Lines link the same patient sample under the different conditions. Mean ± SD shown. * p<0.05, **p<0.001, *** p<0.0001 by Kruskal-Wallis test in (c) and Wilcoxon matched-pair test in (d,e).

To examine how each sorted T cell population contributed to the variance across diseases, we performed principal component (PC) analysis on the genes differentially expressed by ICI-arthritis, RA and PsA T cells (ANOVA with a q-value <0.05). PC1 accounted for 19% of the variance, separating ICI-arthritis T cells from RA and PsA T cells. CD38^hi^ CD127^−^ PD-1^int^ and PD-1^hi^ populations showed the largest spread, suggesting that these populations, rather than the CD38^−^ CD127^+^ PD-1^int/-^ and CD38^−^ CD127^−^ PD-1^int^ KLRG1^+^ populations, were driving the differences (**Fig. 5b**). To explore the extent of gene expression differences among the specific cell subsets, we calculated gene module scores for each population across the diseases (**Supplementary table 4**). Overall, higher IFN scores were observed in all the ICI-arthritis CD8 T cell populations over RA/PsA. Notably, the expanded CD38^hi^ CD127^−^ PD-1^int^ cell population in ICI-arthritis showed a higher expression of activation and dysfunction modules than did the same population in RA/PsA, while maintaining a similarly high expression of cytotoxicity and proliferation modules (**Fig. 5c**). PD-1^hi^ cells in ICI-arthritis also expressed higher activation and dysfunction scores, yet a lower cytotoxicity score, compared to those from RA/PsA.

Taken together, these transcriptomic findings suggested that CD38^hi^ CD127^−^ CD8 T cells in ICI-arthritis are activated and inflammatory and express a strong IFN signature. We hypothesized that IFN may directly contribute to the phenotypic and functional alterations distinguishing ICI-arthritis from RA/PsA. To test this, we treated synovial fluid mononuclear cells from RA and SpA patients with IFN-*β* or IFN-*γ in vitro*. IFN-*β*, but not IFN-*γ*, induced synovial fluid T cells to acquire a CD38^hi^ CD127^−^ phenotype (**Fig. 5d**). These CD38^hi^ CD127^−^ cells expressed a higher level of granzyme B, perforin and Ki67 compared to the CD38^−^ CD127^+^ cells, such that the frequency of CD38^hi^ perforin^+^ cells was also increased by IFN-! (**Fig. 5d, S6b-e**). At the early timepoint, IFN-*β* also increased the total number of perforin^+^ T cells among total CD8 T cells, with no effect on granzyme B or Ki67 expression (**Fig. 5e, S6b-e**). These results support the notion that Type I IFN contributes directly to the accumulation of cytotoxic CD38^hi^ CD127^−^ CD8 T cells in ICI-arthritis.

### Circulating CD38^hi^ CD8 T cells expand in ICI-arthritis

To investigate whether the findings from synovial fluid were translatable to blood, we compared peripheral blood mononuclear cells (PBMC) from individuals with ICI-arthritis to those from non-inflammatory controls, individuals with ICI-associated thyroiditis (ICI-thyroiditis), RA, PsA and systemic lupus erythematosus (SLE), a disease characterized by high IFN production ^(13)^. CD38^hi^ CD127^−^ CD8 T cells were most abundant in blood from SLE patients, a finding consistent with a prior report on SLE ^(24)^. Across the arthritis conditions, the frequency of CD38^hi^ CD127^−^ CD8 T cells was significantly higher in PBMC from ICI-arthritis patients compared to RA and PsA patients. Notably, these cells were not expanded in ICI-thyroiditis patients (**Fig. 6a-b**). The ICI-arthritis and ICI-thyroiditis patients were similar in their ICI-therapy duration and irAE onset time, but differed in age and number of irAEs. Approximately 70% of ICI-arthritis patients studied here experienced irAE involving other tissues and organs such as colitis, hepatitis, pericarditis, skin rash, thyroiditis, peripheral neuropathy, and stroke, while only 40% of the ICI-thyroiditis cohort reported other irAEs, including skin rash and colitis. These results suggest that an increased level of circulating CD38^hi^ CD127^−^ CD8 T cells may be associated with a systemically more aggressive inflammatory state in ICI-treated individuals.

**Figure 6.**
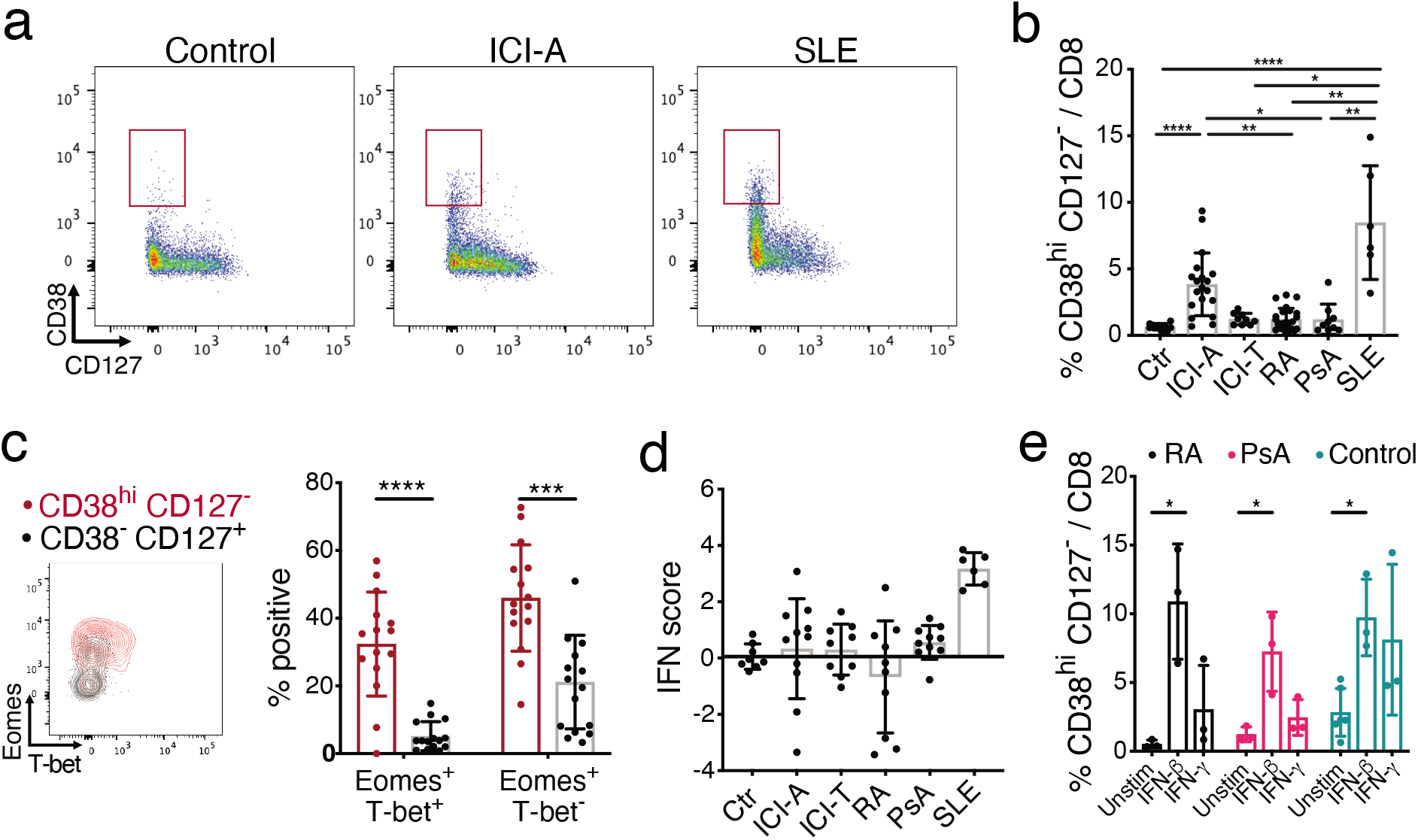
Expanded circulating CD38^hi^ CD127^−^ CD8 T cells in ICI arthritis patients. **a)** Representative flow cytometric plots of CD38 and CD127 expression on gated CD8 T cells from PBMC from non-inflammatory control, ICI-arthritis (ICI-A), and SLE patients. **b)** Frequency of CD38hi CD127-cells among CD8 T cells from PBMC from control (n=10), ICI-A (n=25), ICI-thyroiditis (ICI-T) (n=9), RA (n=22), PsA (n=9), and SLE (n=56) patients. **c)** Representative flow cytometric plot and summarized frequency of intracellular Eomes and T-bet in CD38^hi^ CD127^−^ and CD38^−^ CD127^+^ CD8 T cells from ICI-A PBMC (n=15). **d)** 8 gene-derived IFN score of PBMC from controls (n=8), ICI-A (n=11), ICI-T 11), RA (n=10), PsA (n=10), and SLE (n=6) patients. **e)**Frequency of CD38^hi^ CD127^−^ cells among CD8 T cells from RA, PsA or control PBMC cultured with IFN-β or IFN-γ for 3 days (n=3-5). Mean ± SD shown. * p<0.05, **p<0.001, *** p<0.0001 **** p<0.00001 by Kruskal-Wallis test in (b), (c), (d) and (e).

Consistent with the phenotype of effector cells in synovial fluid, this CD38^hi^ CD127^−^ CD8 T cell population in ICI-arthritis blood showed an enrichment of transcription factors Eomes and Tbet over its CD38^−^ CD127^−^ counterpart (**Fig. 6c**). Unlike with synovial fluid cells, an IFN signature was not consistently detected in PBMC from ICI-arthritis patients, suggesting that IFN may act locally within the joints (**Fig. 6d**). Still, as with synovial fluid cells, IFN-! treatment *in vitro* induced CD8 T cells from PBMCs, from RA, PsA and even controls to acquire a CD38^hi^ CD127^−^ phenotype (**Fig. 6e**). These results support the similarity of surface markers, effector function, proliferating feature and cytokine signaling between CD38^hi^CD127^−^ CD8 T cells in synovial fluid and blood and provide a circulating cell population linked to phenotypically and transcriptionally to a locally enriched T cell subset in ICI-arthritis.

## Discussion

Here we have identified CD38^hi^ CD127^−^ CD8 T cells as a predominant T cell constituent found within the target tissue of the immune adverse condition ICI-arthritis. Cytometric, transcriptomic, TCR repertoire, and functional studies strongly implicate this CD8 T cell subset as a cytotoxic effector population that is activated by ICI therapy and clonally expands within joints to promote synovial inflammation and articular damage.

Efforts to understand T cell roles in autoimmune arthritis have often focused on CD4 T cells, including the PD-1^hi^ Tph cell population in RA joints ^(9, 10)^. We find that ICI-arthritis caused by PD-1 pathway blockade does not induce a large Tph cell population, similar to the low frequency of Tph cells in seronegative arthritides (9). Rather, ICI-arthritis involves marked expansion of a CD38^hi^ CD127^−^ CD8 population, which is infrequent in both RA and PsA. While RA, PsA, and ICI-arthritis have comparably sized synovial CD8 T cell populations ^(10, 11)^, the unique expansion of a CD38^hi^ CD127^−^ subset in ICI-arthritis suggests that the proximal T cell responses differ in these three arthritides and implies that therapies developed to target pathologic T cell responses in RA or PsA may not effectively target the most relevant pathways in ICI-arthritis.

Comparing T cells from RA and PsA highlighted the expression of IFN-inducible genes across ICI-arthritis T cells. Our studies establish Type I IFN as sufficient to induce synovial T cells into a CD38^hi^ perforin^+^ phenotype, implicating a unique role for Type I IFN in influencing T cell responses in ICI-arthritis. This contrasts how Type I IFN has generally not been implicated in PsA pathogenesis and is elevated only modestly in a subset of RA patients ^(25–27)^. Further, while Type I IFN plays a major role in SLE, ICI therapy rarely results in SLE-like disease ^(6)^. Thus ICI-arthritis has a unique immunopathology that does not mirror these paradigmatic spontaneous autoimmune diseases. While blocking Type I IFN signaling, as done in SLE, could be considered as a therapeutic strategy to treat irAEs; ^(28)^ this approach risks compromising the anti-tumor response given the reported positive role of IFNs in ICI-induced anti-tumor T cell activation ^(29–32)^.

The large population of CD38^hi^ CD127^−^ CD8 T cells in ICI-arthritis joints expresses cytotoxic, effector, and proliferative gene programs and produces ample IFN-*γ* and TNF upon restimulation, suggesting a robust inflammatory capacity. These results are consistent with the features of activated CD8 T cells in the gut of patients with colitis after ICI therapy, which also show cytotoxic and proliferative features ^(33)^. Infiltration of CD8 T cells has been noted in ICI-induced immune injury in other organs as well, including liver ^(34)^, skin ^(35)^, heart ^(36, 37)^, and thyroid (38), suggesting that cytotoxic CD8 T cell activation may be a common feature of irAE immune responses, although cytotoxic CD4 T cells have also been implicated ^(39)^.

Our transcriptomic analyses suggest that the T cell infiltrate in ICI-arthritis does not contain a large dysfunctional population, as has been demonstrated in infiltrating lymphocytes in multiple tumor types ^(15, 40–43)^. The CD38^hi^ CD127^−^ population in ICI-arthritis joints does not display classic dysfuncational features such as high PD-1 protein nor transcript levels, and scRNA-seq clearly distinguish the CD38^hi^ CD127^−^ population from a smaller cluster of dysfunctional cells. Further, we found limited clonal overlap between the CD38^hi^ CD127^−^ cluster and the dysfunctional cluster. These results suggest that the predominant CD8 T cells within an active irAE site differ from dysfunctional tumor infiltrating T cells. Notably, CD38^+^ CD8 T cells have been identified in lung tumors ^(44, 45)^, tumors in murine models ^(46, 47)^ and in the blood of patients with SLE. In some cases, these CD38^+^ cells show reduced function or susceptibility to inhibition by adenosine ^(24, 46, 47)^, while in other cases they appear highly functional ^(43)^, as in our analyses of ICI-arthritis. Comparing the functions and TCRs of CD38^hi^ T cells from ICI-arthritis joints with those from tumors, particularly within the same patient, may clarify the functional and clonal relationships of ICI mediated anti-tumor and irAE responses.

Repertoire analyses of T cell in colitis after ICI therapy indicated a relationship between activated T cells and T resident memory cells, consistent with local activation of a resident population ^(33)^. Our analysis of two different joints from the same patient, obtained 6 months apart, revealed multiple shared expanded clones, in particular within the CD38^hi^ CD127^−^ cluster. The presence of common clones in two separate joints indicates that the T cell response in ICI-arthritis can be systemic, rather than a restricted local break in tolerance in a single joint. The detection of expanded clones in the circulation prior to irAEs induced by ipilimumab (anti-CTLA-4) are consistent with this idea that autoreactive clones are present systemically ^(48)^. It will be of major interest to determine whether T cells in different irAE sites in the same patient express the same TCRs, suggesting targeting of shared antigens in different tissues.

Consistent with a systemic response, we detected an increased frequency of CD38^hi^ CD127^−^ CD8 T cells in the circulation of patients with ICI-arthritis. We did not similarly detect a consistent increase in IFN-induced genes in PBMC, suggesting that IFNs may act locally within tissues, in contrast to the systemic IFN signature in SLE ^(49, 50)^. Expansion of CD38^+^ cells is consistent with prior observations that in the peripheral blood of melanoma patients, ICI therapy increases the frequency of Ki67^+^ CD38^+^ HLA-DR^+^ cells, which show clonal overlap with tumor infiltrating lymphocytes ^(47, 51)^. Notably, we did not detect an increase in CD38^hi^ CD127^−^ cells in a cohort of patients with ICI-thyroiditis, consistent with recent findings emphasizing CD4 T cell alterations in ICI-thyroiditis ^(39)^. This suggests that PD-1 blockade does not universally expand CD38^hi^ CD127^−^ CD8 cells in all patients. The ICI-arthritis cohort had higher frequency of other irAEs compared to the ICI-thyroiditis cohort; thus, we hypothesize that expansion of CD38^hi^ cells reflects the extent of T cell activation after PD-1 blockade. Determining whether circulating CD38^hi^ cells share TCRs with those within irAE target tissue will be of interest, which if also found prior to ICI therapy in the blood, could be studied as biomarkers for irAEs. Further, evaluating the extent to which circulating CD38^hi^ cells associate with specific irAEs and their relationship to anti-tumor responses will be important to explore with longitudinal study of a larger cohort of patients.

Our study is limited by relatively small number of patients studied, by the lack of paired tumor and blood samples, and by the absence of therapeutic response data, which we expect will be spurred by the findings described here. In sum, our study defines characteristic differences between the T cell response in ICI-arthritis and that in two common forms of spontaneous arthritis. These comparative analyses raise caution about the abilty to coopt treatment paradigms from spontaneous autoimmunity to treat irAEs and here nominate an IFN-induced CD38^hi^ CD127^−^ CD8 cells as a likely pathologic driver of irAEs following ICI therapy that may be targetable therapeutically.

## Methods

### Human subjects research

Human subjects research was performed according to the Institutional Review Boards at Mass General Brigham (IRB protocol 2014P002558) or Hospital for Special Surgery (IRB protocols 2017-1898 and 2014-233) via approved protocols with informed consent as required. Synovial fluid samples were collected from patients with ICI-arthritis, RA or PsA as discarded fluid from clinically-indicated arthrocenteses. Patients with ICI-arthritis developed arthritis after receiving checkpoint inhibitor therapy to treat a malignancy and were diagnosed by experienced rheumatologists. Type and status of cancer, type and duration of CI therapy, tender or swollen joint counts (mono-, oligo- or poly- arthritis), other irAE, prior history of rheumatologic conditions, serum rheumatoid factor (RF) and anti-CCP antibody status, C-reactive protein level, and medication usage were obtained by review of electronic medical records. Seropositive (RF+ and/or anti-CCP+) RA patients fulfilled 2010 ACR/EULAR classification criteria. PsA patients were diagnosed with PsA by their treating rheumatologist. Blood samples were obtained from individuals with ICI-arthritis, ICI-thyroiditis, seropositive RA, PsA, SLE as well as individuals without inflammatory arthritis. Mononuclear cells from synovial fluid and peripheral blood were isolated by density centrifugation using Ficoll-Paque Plus (GE healthcare) and cryopreserved in FBS + 10% DMSO by slow freeze, followed by storage in liquid nitrogen for batched analyses. For experimental analyses, cryopreserved samples were thawed into RPMI medium + 10% FBS.

### Mass cytometry staining

Cryopreserved synovial fluid cells were thawed and trypan blue negative viable cells were counted by hemocytometry. Approximately 1 million live cells per sample were used for mass cytometry staining. All antibodies were obtained from the Longwood Medical Area CyTOF Core except for anti-PD-1 antibody (MIH4), which was conjugated with heavy-metal isotope (Fluidigm kit) and validated in house. Buffers were from Fluidigm unless otherwise specified. Cells were stained with cisplatin (Fluidigm) for viability then washed. Surface antibody cocktail was prepared in cell staining buffer (Fluidigm) and added to all samples equally after Fc block (BD). Cells were washed then fixed and permeabilized using eBioscience Transcription Factor Fix/Perm Buffer followed by barcoding (Fluidigm). Barcoded samples were pooled together and stained with the intracellular antibody cocktail in intracellular staining buffer (Fluidigm). Cells were re-fixed in 4% formalin (Sigma-Aldrich). Intercalator-Ir was diluted in CyTOF PBS and applied to cells. Cells were then washed and resuspended with cell acquisition solution (Fluidigm) containing 1:10 diluted EQ beads (Fluidigm). Acquisition was performed on a CyTOF-Helios mass cytometer (Fluidigm).

### Mass cytometry data analysis

Cytometry data were normalized and debarcoded as previously described ^(52)^. Live cells (DNA^+^ 195_Pt^−^ 140_Beads^−^) were first gated prior to gating for specific cell populations using the following scheme: monocytes (CD3^−^ CD14^+^), B cells (CD14^−^ CD19^+^), and T cells (CD3^+^ CD14^−^). Gated populations from 6 ICI-arthritis, 5 RA and 5 SpA were concatenated for high-dimensional analyses using the implementations on Cytobank (www.cytobank.org). For donors with more than 10,000 cells, we randomly selected 10,000 cells to ensure that samples were equally represented. In this way, we created downsampled datasets of 160,000 viable cells or 42,144 CD8^+^ T cells from 16 samples for analysis. Dimensional reduction was performed using viSNE algorithm ^(53)^ using 2000 iterations with a perplexity = 30 and a theta = 0.5. Hierarchical consensus clustering was performed using FlowSOM algorithm to generate 15 metaclusters and 225 clusters using 100 iterations ^(54)^. Antibody channels excluding gating markers were used for analyses. Heatmaps of row-normalized median expression of representing markers in the metaclusters are shown, in which the metaclusters were arranged by hierarchical consensus clustering. Metaclusters and markers of interest were overlaid on tSNE plot for visualization. Manual biaxial gating was performed using FlowJo v.10.4.2 for quality control and independent examination of the expression of markers and frequencies of populations.

### Flow cytometry staining

Cryopreserved cells were thawed, washed and counted. Cells were stained in PBS with Aqua fixable live/dead dye (Invitrogen) for 15 minutes at room temperature and washed. For surface staining, cells were then stained in PBS with 1% BSA with the following antibodies for 30 minutes at 4 °C: anti-CD14-BV510-dump (M5E2), anti-CD25-FITC (M-A251), anti-CD8-BUV395 (RPA-T8), anti-CD4-BV605 (RPA-T4), anti-CD38-PercpCy5.5 (HIT2), anti-CD45RA-BV711 (HI100), anti-PD-1-PE-Cy7 (MIH4), anti-CD3-AF700 (UCHT1), anti-CD127-APC (A019D5) and anti-KLRG1-BV421 (SA231A2) from BioLegend. Cells were washed in cold PBS, passed through a 70-micron filter, and data acquired on a BD Fortessa analyzer using FACSDiva software. Data were analyzed using FlowJo 10.4.2. For intracellular staining of transcription factors, cells were processed and stained for viability and indicated surface markers as described above. Cells were washed and incubated with 1x eBioscience Transcription Factor Fixation/Permeabilization Buffer at room temperature for 1 hour. Cells were then washed in 1x eBioscience Permeabilization Buffer twice and incubated with indicated intracellular antibodies including anti-T-bet-BV785 (4B10, BD Biosciences), anti-EOMES-PerCP-ef710 (X4-83, BD Biosciences), anti-granzyme B-AF647 (GB11, Invitrogen), anti-granzyme K-FITC (GM26E7, Invitrogen), anti-perforin-BV421 (dG9, Invitrogen) and anti-Ki-67-BV605 (Ki67, BioLegend) at room temperature for 1 hour. Cells were washed twice in 1x eBioscience Permeabilization Buffer, passed through a 70-micron filter, and data acquired on a BD Fortessa analyzer. For intracellular detection of TNF and IFN-*γ*, sorted cells were stimulated with 1x PMA/ionomycin + Brefeldin A/Monesin (Invitrogen) at 37 °C for 2 hours. Cells were stained for viability and indicated surface markers as described above. Cells were washed and incubated with 1x eBioscience Transcription Factor Fixation/Permeabilization Buffer at room temperature for 1 hour. Cells were then washed in 1x eBioscience Permeabilization Buffer twice and incubated with indicated intracellular antibodies including anti-TNF-PE (MAb1) and anti-IFN-*γ*-APC (4S.B3) from BioLegend at room temperature for 1 hour. Cells were washed twice in 1x eBioscience Permeabilization Buffer, passed through a 70-micron filter, and data acquired on a BD Fortessa analyzer.

### Flow cytometric cell sorting

An 8-color flow cytometry panel was developed to identify CD8 T cell populations within SFMC. Antibodies include anti-CD14-BV510 (M5E2), anti-3-Alexa Fluor 700 (UCHT1), anti-CD8-BV510 (RPA-T8), anti-PD-1-PE-Cy7 (MIH4), anti-CD127-APC (A019D5), anti-CD38-PE (HIT2), anti-KLRG1-BV421 (SA231A2) and propidium iodide (all from BioLegend). SFMC were incubated at 4 °C with antibodies in HBSS/1% BSA for 30 min. Cells were washed once in HBSS/1% BSA, centrifuged and passed through a 70-μm filter and propidium iodide was added immediately prior to sorting. Cells were sorted on a 4-laser BD FACSAria Fusion cell sorter. Intact cells were gated according to forward scatter and side scatter area (FSC-A and SSC-A).

Doublets were excluded by serial FSC-H/FSC-W and SSC-H/SSC-W gates (H, height; W, width). Non-viable cells were excluded based on propidium iodide uptake. Cells were sorted through a 70-μm nozzle at 70 psi. Cell purity was routinely >98%. For functional analyses, approximately 200,000 cells were sorted from each population into cold RPMI/10% FBS. For RNA-seq, up to 1,500 cells were collected from each cell subset directly into buffer TCL (Qiagen) with 1% β-mercaptoethanol (Sigma). Flow cytometric quantification of cell populations was performed using FlowJo v.10.0.7.

### Sample preparation for low-input bulk RNA-seq

RNA was isolated from 1,500 cells from sorted T cell subpopulations. 5uL of total RNA were placed in wells of a 96-well plate and RNA-seq libraries were prepared at Broad Technology Labs at the Broad Institute of Harvard and MIT using the Illumina SmartSeq2 platform. Libraries were sequenced to generate 38 base paired-end reads.

### Low-input bulk RNA-seq analysis

Sequencing samples were examined with FastQC for quality control and trimmed with trimmomatic. Samples with low sequencing quality were removed and 86 samples were used for downstream analysis (**Fig. S3a**). Reads were mapped to GRCh38 (Ensembl release 96) using STAR alignment software. Lowly expressed genes (log2 FPKM<2) were filtered, and the expression of 17,779 genes were used for downstream analysis (**Fig. S3b**). TMM normalized read counts were generated and data was log2 transformed, batched corrected and scaling was applied (mean=0, variation=1) when appropriate. Differential expression analysis and gene set enrichment analysis was performed in Qlucore software. In comparisons between two specific cell populations by t-test, genes with log fold change >1.5 and FDR <5% were considered differentially expressed. In comparisons among more than two cell populations by ANOVA, genes with an FDR <1% were considered differentially expressed.

### Analysis of gene module score and IFN response score

Module scores based on specific gene sets (memory, effector, proliferation, IFN-inducible genes etc.) were calculated for each population as the average of the scaled (Z-normalized) expression of the genes in the list. A similar approach was used when calculating gene set-based scores in bulk RNA-seq data.

Module gene lists were compiled from published studies and molecular signature databases. (MSigDB v7.4 from UC San Diego and Broad Institute)

Memory module genes (Tcm, Tem, Trm and Temra) were from Zhang et al, Nature 2018^(14)^. Dysfunction module genes were from Zhang et al, Nature 2018^(14)^ and Li et al, Cell 2019^(15)^. Cytotoxicity module genes were from Li et al, Cell 2019^(15)^, and MSigDB gene sets M16355 (BIOCARTA_NK cells pathway), M13247 (BIOCARTA_ T Cytotoxic Cell Surface Molecules) and M5669 (KEGG_ Natural killer cell mediated cytotoxicity) Proliferation module genes were from Gene Ontology gene sets (GO_G1_S_TRANSITION_OF_MITOTIC_CELL_CYCLE, GO_G2_M_TRANSITION_OF_MITOTIC_CELL_CYCLE) and Reactome MSigDB gene sets M1017 (DNA_REPLICATION), M1080 (G2_M_DNA_DAMAGE_CHECKPOINT), M27662 (M_PHASE), M17283 (MITOTIC_G1_PHASE_AND_G1_S_TRANSITION), M27673 (MITOTIC_SPINDLE_CHECKPOINT) and M3158 (S_PHASE) Effector module genes were from MSigDB gene sets M3044 (GOLDRATH_EFF_VS_MEMORY_CD8_TCELL_DN), M3041 (GOLDRATH_EFF_VS_MEMORY_CD8_TCELL_UP), M5834 (GSE9650_EFFECTOR_VS_EXHAUSTED_CD8_TCELL_DN), M5837 (GSE9650_EFFECTOR_VS_MEMORY_CD8_TCELL_DN), M3073 (GSE10239_MEMORY_VS_DAY4.5_EFF_CD8_TCELL_UP), M4407 (GSE22886_NAIVE_CD8_TCELL_VS_MEMORY_TCELL_DN), M8435 (GSE23321_CENTRAL_MEMORY_VS_NAIVE_CD8_TCELL_UP), M9490 (GSE41867_DAY6_EFFECTOR_VS_DAY30_EXHAUSTED_CD8_TCELL_LCMV_CLONE 13_DN), M9492 (GSE41867_DAY8_EFFECTOR_VS_DAY30_EXHAUSTED_CD8_TCELL_LCMV_CLONE 13_DN), M9480 (GSE41867_MEMORY_VS_EXHAUSTED_CD8_TCELL_DAY30_LCMV_UP) and M3027 (KAECH_DAY8_EFF_VS_MEMORY_CD8_TCELL_UP) IFN-inducible genes were from Arazi, et al, Nat Immunol, 2019^(20)^. The compiling processes were based on 2 principles: 1). integration: the collection of genes from gene sets of the same module were merged; 2). discrimination: genes shared by memory-effector-dysfunction categories were excluded from memory and dysfunction module and considered effector, (as an intermediate state between memory-dysfunction) while genes shared by cytotoxicity, proliferation or IFN-inducible genes were not excluded from other modules. Differentially expressed module genes were obtained from the intersection of each of the above module genes and the ANOVA-tested, batch-corrected DEGs (q < 0.01) among the sorted bulk populations (**Supplementary table 3**).

### Sample preparation for single cell RNA-seq

#### Synovial fluid

Mononuclear cells from synovial fluid were isolated using Ficoll-Paque Plus (GE healthcare) and cryopreserved in Cryostor10 (BioLife) in liquid nitrogen. For experimental analyses, cryopreserved samples were thawed into RPMI medium (Corning) + 10% FBS (HyClone) + 1% L-glutamine (Gibco), referred to as RPMI++.

#### Synovial tissue

Fresh synovial tissues were cut into 3-mm^3^ fragments and preserved in Cryostor (BioLife). For experimental processing, tissues were thawed into RPMI++. Following two subsequent washes in RPMI, the tissues were finely chopped and transferred into 5ml polystyrene tubes () containing digestion buffer (5ml/sample) made with RPMI medium + Liberase TL (100μg/ml; Roche) + DNaseI (100μg/ml; Roche). The tubes were placed securely in a MACSmix tube rotator (Miltenyi Biotec) and placed in an incubator at 37ºC, 5% CO2 for 30 minutes. The digested tissue was filtered over 70μM cell strainer (BD) and subjected to further mechanical dissociation using a syringe plunger. The eluate was washed with RMPI++ and centrifuged at 1500 rpm for 4 minutes at 4ºC. The cells were resuspended in RPMI++ and passed through a second filtration using 40μM cell strainer (BD). Cells were then counted and used for downstream applications.

Single cell suspensions from fluid and tissue were subjected to Human TruStain FcX (Biolegend) in FACS buffer (PBS supplemented with 5% FBS (HyClone)) prior to staining. The antibodies used for identification of CD8 T-cells were anti-CD3 (APC, clone UCHT1, Biolegend) and CD8 (PerCP/Cyanine5.5, clone RPA-T8, Biolegend). Cells were sorted on a three-laser BD FACSAria Fusion cell sorter at the Flow Cytometry Core Facility at Weill Cornell Medicine (WCM). Intact cells were gated according to forward scatter and side scatter area (FSC-A and SSC-A). Doublets were excluded by serial FSC-H/FSC-W and SSC-H/SSC-W gates (H, height; W, width). Non-viable cells were excluded based on DAPI uptake. Cells were further selected for CD3 and CD8 surface expression and sorted through a 70-μm nozzle at 70 psi.

### Single cell RNA-seq library preparation

5’ gene expression (GEX) and paired TCR libraries for single cell RNA-seq were prepared using the 5’ Chromium Next GEM Single Cell v2 (Dual Index) reagents and protocol provided by 10X Genomics. The pooled 5’ GEX and TCR libraries at 10nM concentration were sequenced using NovaSeq6000 S1 Flow Cell by the Illumina platform at Genomics Research Core Facility at WCM.

### scRNA-seq data analysis

scRNA-seq data processing and alignment: 10x FASTQ files were processed with the Cellranger count 4.0 pipeline with default parameters. Reads were aligned to the human reference sequence GRCh38. Seurat package (v.4.0.0) was used to perform unbiased clustering of the CD8^+^ sorted T cells from our patient samples. QC was performed to remove cells that had less than 50 genes, more than 3000 genes, or >10% mitochondrial gene expression, resulting in a total of 18,472 cells and 19,406 genes. The dataset was then log-normalized using a scale factor of 10,000. Potential confounders such as percent mitochondrial gene expression and number of UMI per cell was regressed out during scaling (mean of 0 and variance of 1) and before principal component analysis, which used the top 2000 highly variable genes. Elbow plot was used to determine the statistically significant principal components, where then we used the first 21 PCs for follow-up analysis. Harmony (v1.0) was performed to improve integration and correct for batch effects on our samples, with parameters of max.iter.cluster = 30, and max.iter.harmony = 20 and sample as the only covariate. Eight clusters were found at 0.5 resolution, and their identity were annotated based on the expression of differentially expressed genes (DEG) using FindAllMarkers function using default parameters, where only genes detected in at least 25% of the cells in the two comparison sets are used. The AddModuleScore function in Seurat was used to caluculate module scores, where the average expression level of genes of interest are subtracted from a randomly aggregrated expression of control features, to distinguish clusters.

### TCR analyses

Cellranger vdj 4.0 pipeline was used to generate clonotype information from 10x 5’ VDJ FASTQ files, with default parameters. Reads were aligned to the human reference sequence GRCh38. Clonotype information was then manually inputted into our Seurat object as metadata information, using the cell barcodes to match the clonotype information to our cells. The integration resulted in 6843 unique clonotypes, where all cells within a clonotype share the same CDR3 alpha and CDR3 beta sequences. This allowed us to explore the relationship between TCR sequence and its phenotype.

### Measures of diversity

#### Simpson’s diversity index

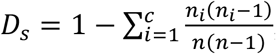; where *n*_*i*_ is the number of cells within the *i*th clonotype, c is the total number of unique clonotypes in our cluster, and n is the total number of cells with clonotypes within our cluster. Simpson Diversity Index ranges from 0 to 1, with 0 being a community with no diversity and 1 being a community of the highest diversity.

#### Shannon Equitability

E = H / S; where H is the Shannon’s diversity index value and S is the number of clonotypes in that sample. Shannon Equitability ranges from 0 to 100 and is based on the Shannon Diversity Index but takes into account the actual number of unique clonotypes present within each cluster to assign a value that can be directly compared across all clusters. As the value approaches 100, the community is approaching maximum diversity.

### RNA extraction, reverse transcription and real-time PCR (qPCR)

RNA isolated using RNeasy Micro Kits (Qiagen). cDNA was prepared using Quantitect RT-PCR (Qiagen) and PCR performed with Brilliant III SYBRGreen on a Stratagene Mx3000. Primers used were as follows: 18S forward: GGGAGCCTGAGAAACGGC, reverse: GGGTCGGGAGTGGGTAATTT. IFI27 forward: GAATCGCCTCGTCCTCCATAG, reverse: CGCCAGGATACTTACCCAGTG. IFI44 forward: CTGAGACGAATGCTATGGGCT, reverse: GACAGAGAGCTGCCAGGTATT. IFI44L forward: CAGATTTGGAACTGGACCCCA, reverse: AGGGCCAGATTACCAGTTTCC. ISG15 forward: CGCAGATCACCCAGAAGATCG, reverse: TTCGTCGCATTTGTCCACCA. IFIT1 forward: TTGATGACGATGAAATGCCTGA, reverse: CAGGTCACCAGACTCCTCAC. IFIT3 forward: TCAGAAGTCTAGTCACTTGGGG, reverse: ACACCTTCGCCCTTTCATTTC. IRF7 forward: CCCACGCTATACCATCTACCT, reverse: GATGTCGTCATAGAGGCTGTTG. RASD2 forward: GAGCGCCACAAAGAAGTGTC, reverse: ACAATGTGTGGGGTCCTTGG.

### Cell culture with IFN stimulation

Cryopreserved mononuclear cells from synovial fluid or peripheral blood were thawed and counted as usual. Cells were plated at 2.5×10^6 cells/mL in 1mL per well in a 12-well plate and cultured with or without IFN-*β* (1kU/mL) or IFN-*γ* (50ng/mL) for 18 hours or 72 hours. At the time of harvest, cells were washed and stained with surface markers and intracellular markers as described above.

### Statistics

Statistical analysis was performed as described in each section and figure legends using Prism 8 software. Unless otherwise stated, data are presented as mean ± SD from data obtained from at least two independent experiments. Parametric and non-parametric analyses were used where appropriate based on testing for a normal distribution using the D’Agostino–Pearson normality test or Shapiro-Wilk normality test. Two-tailed Student’s t test was used for two-group comparisons (Mann-Whitney test was used for nonparametric data). One-way analysis of variation (ANOVA) followed by the Holm-Sidak test was used for multiple comparisons (Kruskal-Wallis test was used for nonparametric data). P-values < 0.05 were considered significant after adjusting for multiple testing where appropriate.

### Data Availability

Bulk and single cell RNA-seq data will be shared through Immport upon publication.

## Acknowledgements

This work has been supported in part by funding from the Rheumatology Research Foundation (to DAR, LTD, ARB), Burroughs Wellcome Fund Career Award in Medical Sciences, NIAMS K08 AR072791, P30 AR070253 (to DAR), NIAMS R01 (to MBB), NIAID R01 AI148435 (LTD). We thank Adam Chicoine and the BWH Human Immunology Center Flow Cytometry Core for cell sorting assistance.

## Author contribution

R. Wang, L. Donlin, A.R. Bass, and D.A. Rao conceived the overall project. K.K. Chan, A.R. Bass, A. Cunningham-Bussel, L. Chen, D.J. Todd, L. MacFarlane, E.M. Massarotti, J.A. Sparks, O.R. Hamnvik, L. Min and A. Tirpack collected human subject data and helped analyze clinical data. R. Wang, K. Marks, and A.H. Jonsson, performed mass cytometry and flow cytometry phenotyping analyses. R. Wang generated and analyzed bulk RNA-seq data, and R. Wang and G. Dunlap analyzed bulk RNA-seq data. K. Marks performed and analyzed cell culture experiments. A. Singaraju and L. Shakib generated scRNA-seq and TCR data, and A. Singaraju, L. Shakib, and G. Dunlap analyzed scRNA-seq/TCR data. M.R. Fein and M.B. Brenner provided experimental advice and assisted with interpretation of T cell data. L.T. Donlin, R. Wang, and D.A. Rao wrote the initial manuscript, and all authors participated in revising the manuscript.

## List of Supplementary Tables

**Supplementary table 1**. Clinical characteristics of ICI arthritis patients with samples studied.

**Supplementary table 2.** Mass cytometry panel for analysis of synovial fluid cells.

**Supplementary table 3.** Genes differentially expressed across CD8 T cell populations from ICI-arthritis samples.

**Supplementary table 4.** Genes used for calculation of module scores.

**Supplementary table 5.** GSEA pathways altered in CD38^hi^ CD127^−^ cells and CD38^−^ CD127^+^ cells.

**Supplementary table 6**. Top genes correlated with CD38, IL7R, and PDCD1 in bulk RNA-seq data.

**Supplementary table 7.** Signature genes for indicated CD8 T cell populations based on differential gene expression in bulk RNA-seq data

**Supplementary table 8.** Reactome pathways differentially expressed between ICI-arthritis T cells and RA+PsA T cells.

**Supplementary table 9.** Genes differentially expressed between ICI-arthritis T cells and RA+PsA T cells.

---

**Supplementary figure 1.**
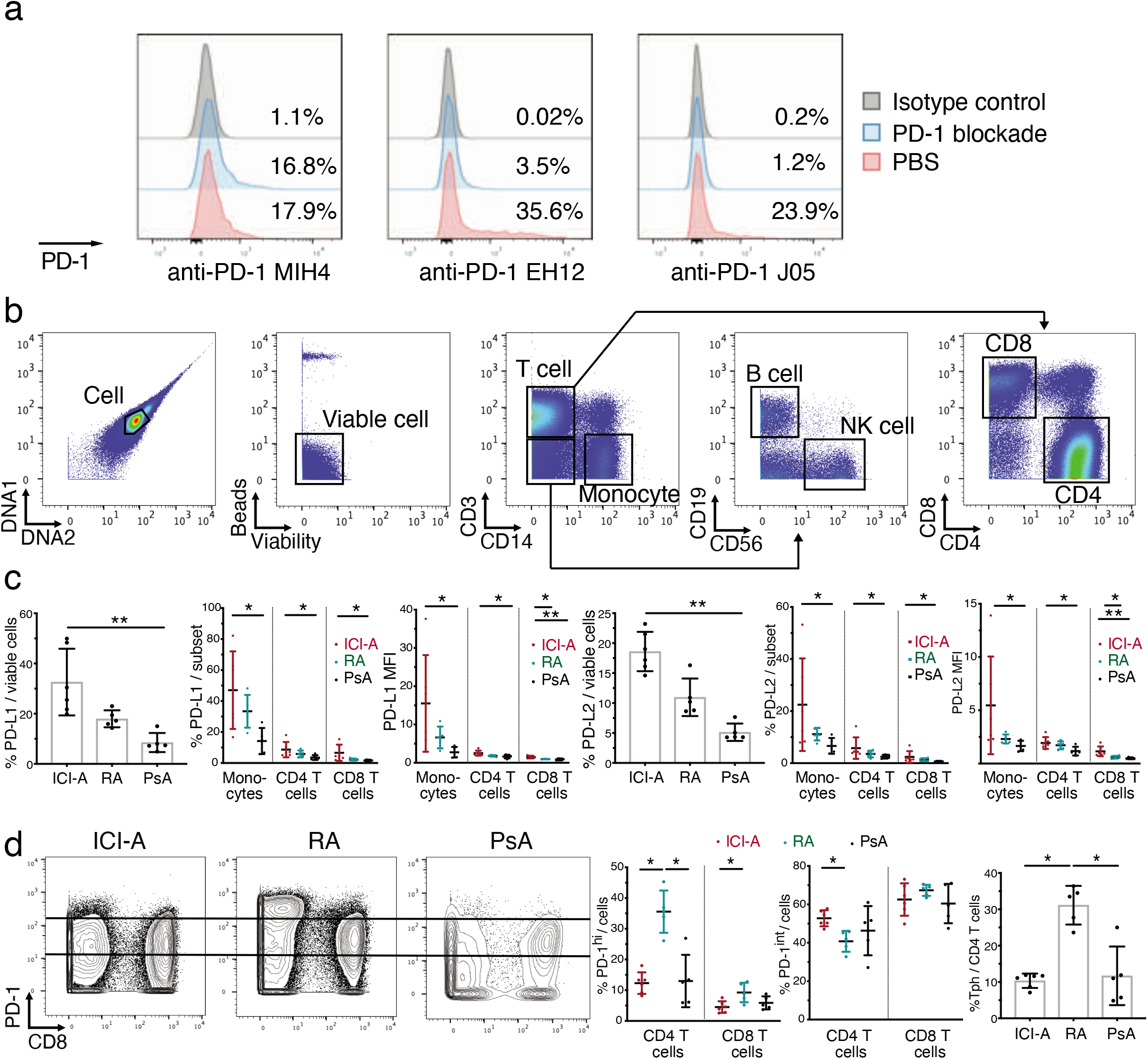
Basic analysis of mass cytometry of ICI-A, RA and PsA synovial fluid. **a)** Flow cytometric detection of PD-1 on T cells using indicated detection antibody clones with or without prior incubation with pembrolizumab blockade in vitro. **b)** Example gating of mass cytometry data. **c)** PD-L1 and PD-L2 expression by percentage and mean fluorescence intensity (MFI) on indicated cell types from ICI-A (n=6), RA (n=5) and PsA (n=5) synovial fluid detected by mass cytometry. **d)** Example gating of high, intermediate and low PD-1 expression levels on T cells from ICI-A, RA and PsA synovial fluid detected by mass cytometry. Quantification of PD-1^hi^ cells, PD-1^int^ cells and PD-1^hi^CXCR5^−^ CD4 Tph cells are shown. Mean ± SD shown. * p<0.05, **p<0.001 by Kruskal-Wallis test in (c) and (d).

**Supplementary figure 2.**
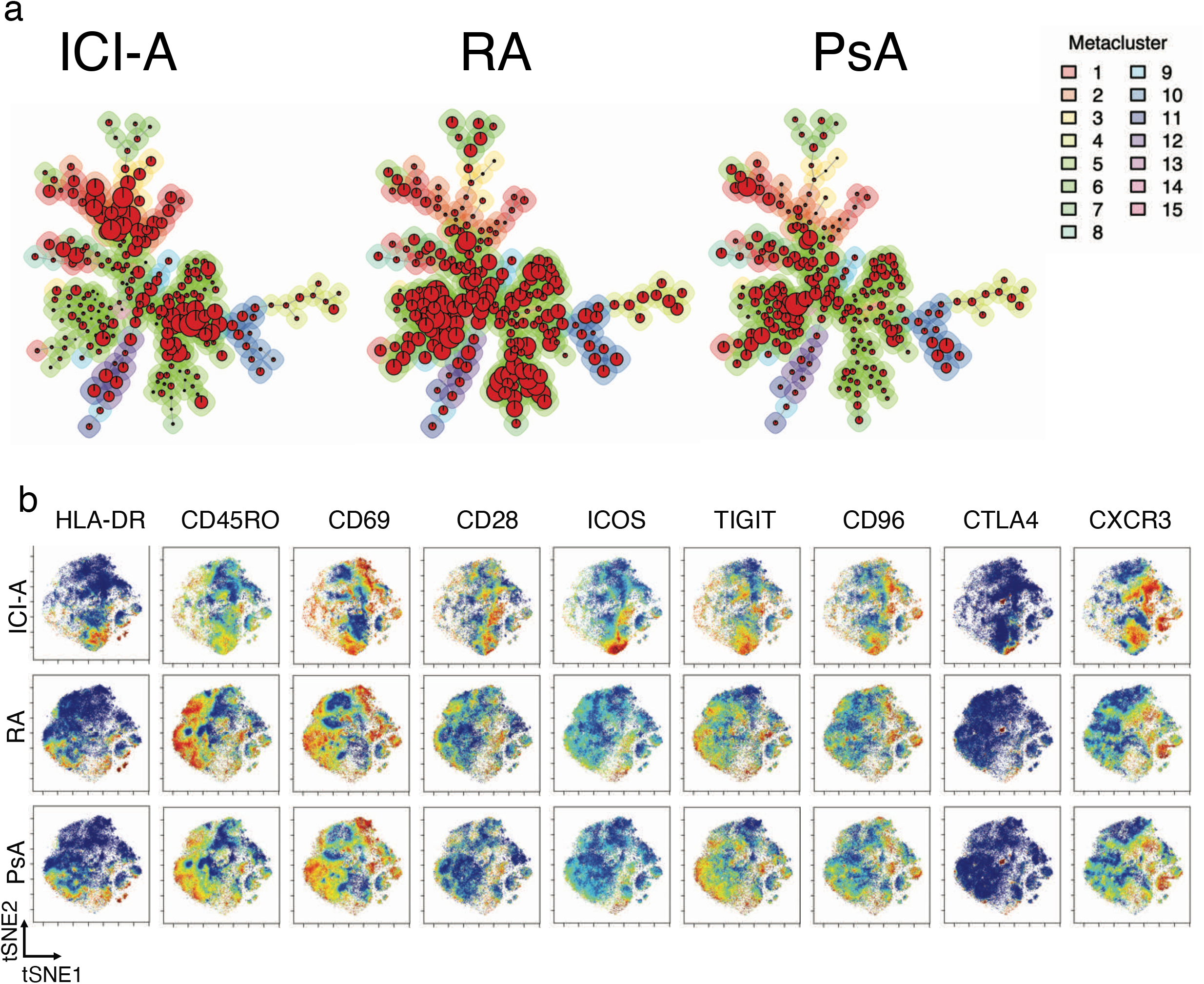
Unsupervised analysis of mass cytometry of ICI-A, RA and PsA synovial fluid. **a)** FlowSOM minimus spanning tree showing metaclusters on CD8 T cells from ICI-A, RA and PsA synovial fluid detected by mass cytometry. **b)** tSNE plots showing expression of indicated markers on CD8 T cells from ICI-A, RA and PsA synovial fluid detected by mass cytometry.

**Supplementary figure 3.**
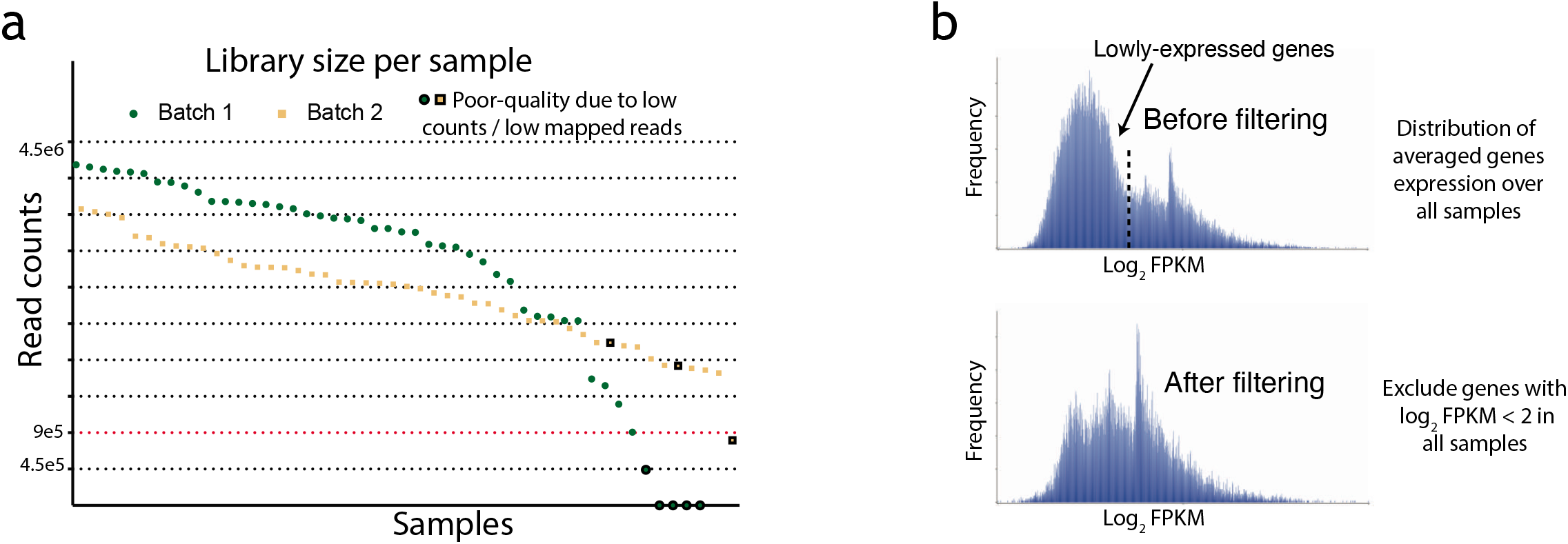
Quality of bulk RNA sequencing data. **a)** Distribution of read counts per sample colored by batch. 6 samples with less than 9×105 reads and 2 samples with poor unique mapping to reference genome were removed. **b)** Distribution of averaged gene expression over all samples before and after filtering.

**Supplementary figure 4.**
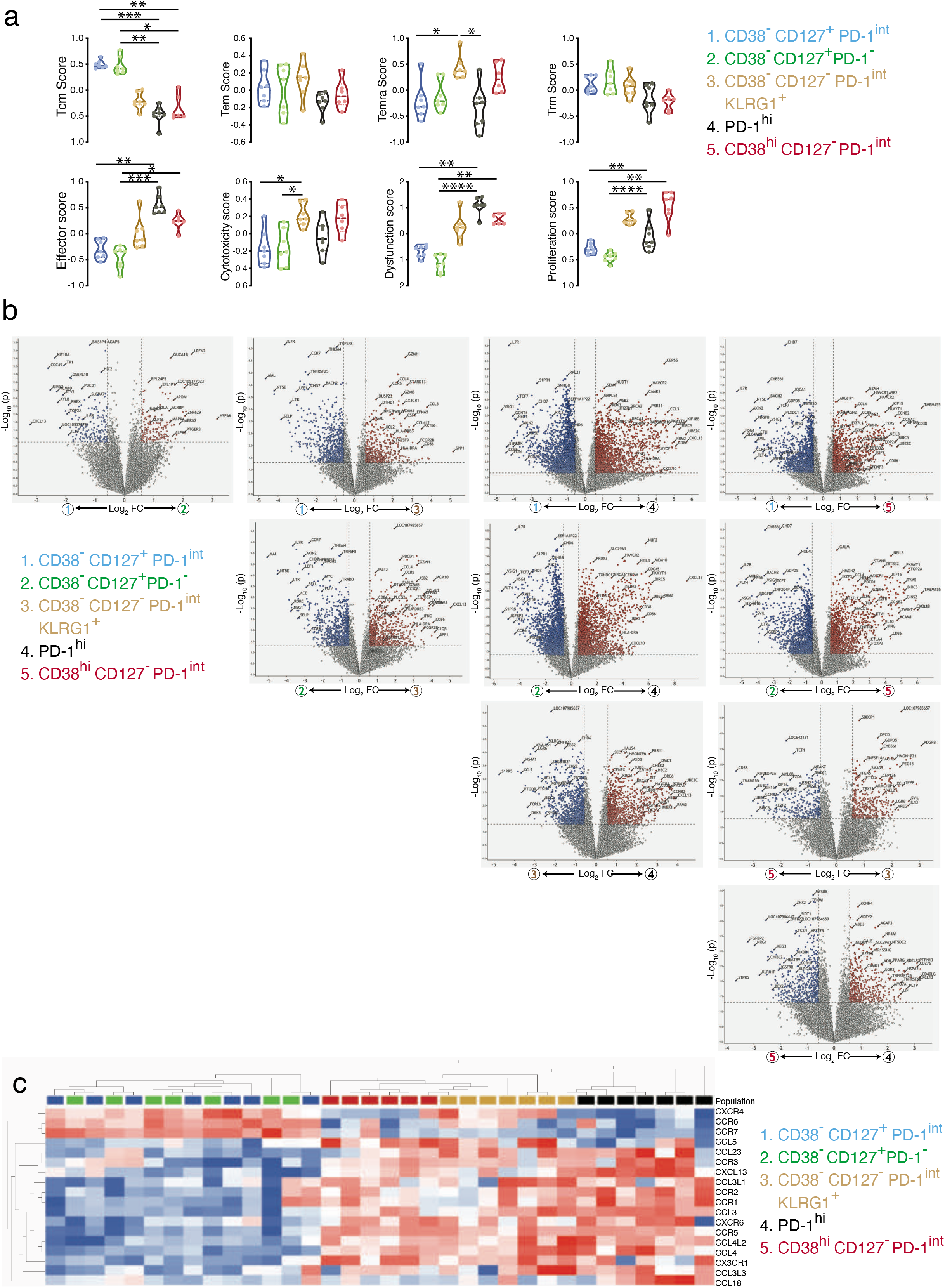
Transcriptomic distinctions between CD8 T populations in ICI-arthritis synovial fluid. **a)** Gene module scores of sorted CD8 T cell populations from ICI-A synovial fluid, calculated based on total module genes. **b)** Volcano plots showing genes differentially expressed by each pair of the 5 sorted CD8 T cell populations from ICI-A. Number of DEGs (q<0.05) are summarized. **c)** Hierarchical clustering of CD8 T cell populations from ICI-A synovial fluid based on row-normalized mean expression of differentially expressed chemokines and chemokine receptors measured by RNA-seq. Mean ± SD shown. * p<0.05, **p<0.001, *** p<0.0001 **** p<0.00001 by Kruskal-Wallis test in (a) and (b).

**Supplementary figure 5.**
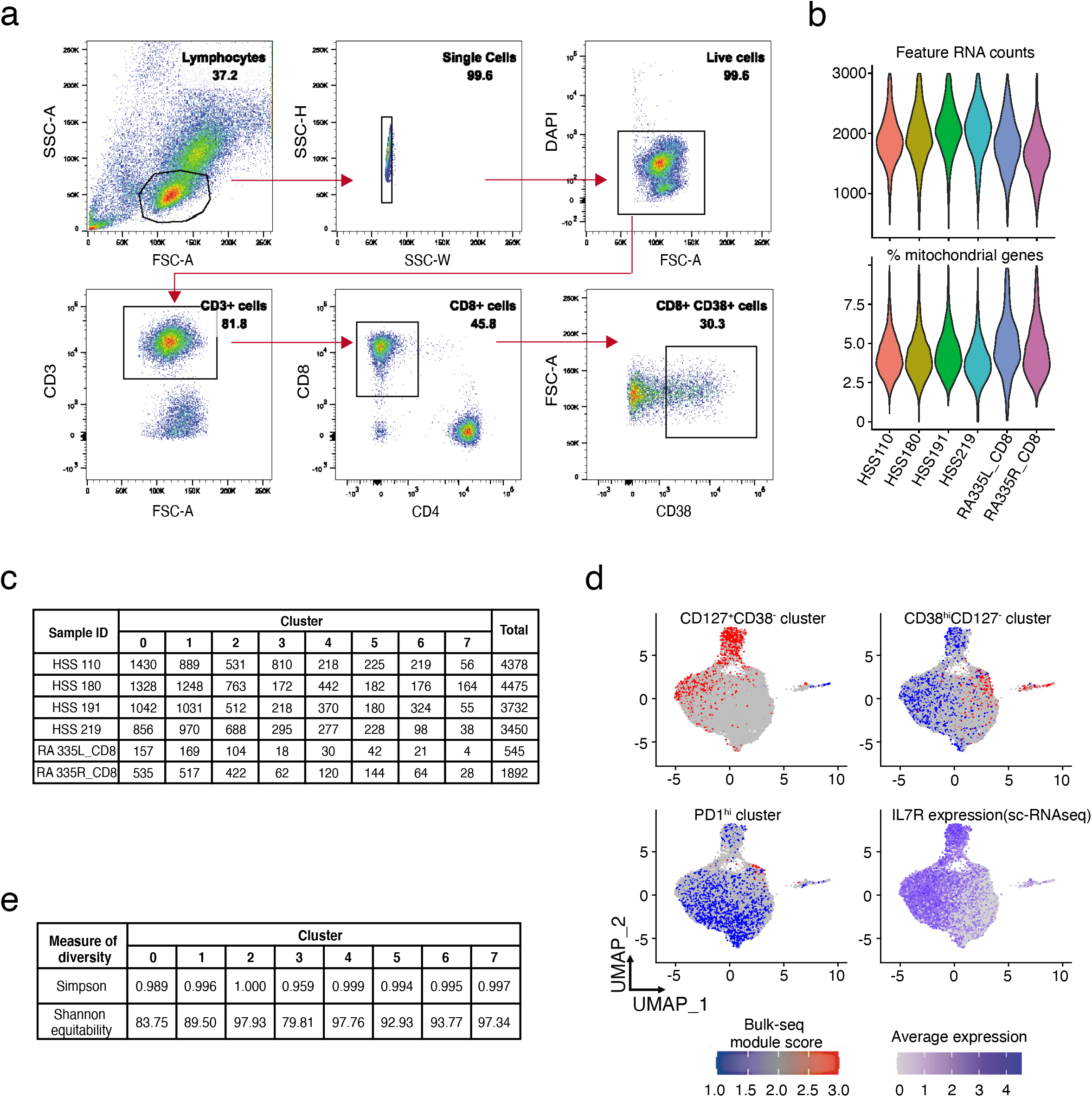
Single-cell analysis of CD8 T cells from ICI-arthritis synovial fluid and tissue. **(a)** Gating scheme of CD8 T cell isolation from ICI-arthritis synovium for single cell RNA sequencing. **b)** Distribution of cells that passed quality control having <3,000 total genes (feature RNA counts) and <10% mitochondrial genes. **c)** Tabulation of QC filtered CD8 T cell numbers contributing to 8 transcriptomically distinct clusters from every individual patient. **d)** Overlay of signature scores for gene sets expressed in CD38^hi^ CD127^−^, CD38^−^ CD127^+^ and PD-1^hi^ populations from bulk RNA-sequencing and UMAP visualization of *IL7R* (CD127) expression across synovial CD8 T cells. **e)** Simpson and Shannon equitability indices calculated for the eight CD8 T cell clusters.

**Supplementary figure 6.**
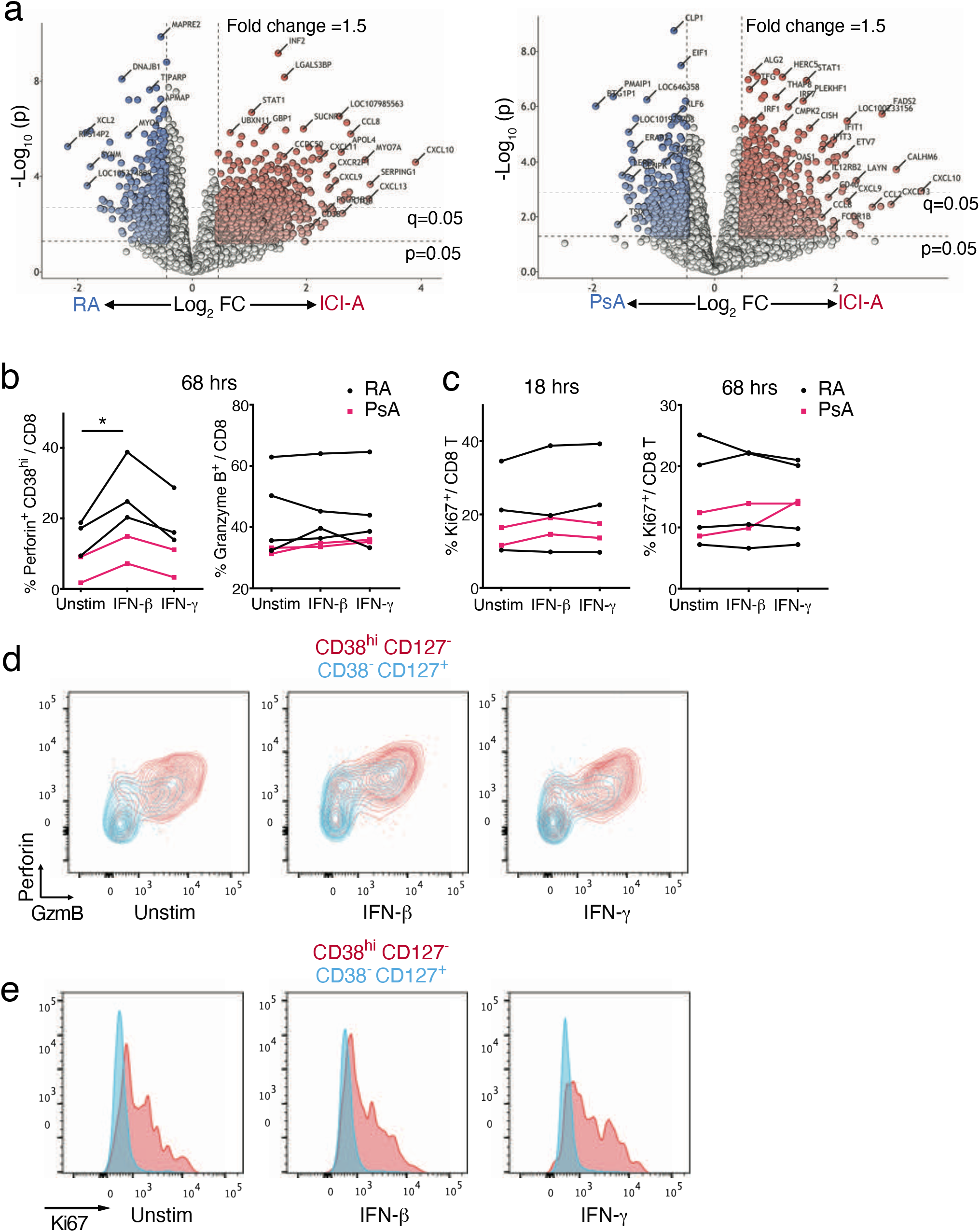
Expression of intracellular markers in RA and PsA SFMC with or without IFN treatment. **a)** Volcano plots showing genes differentially expressed by CD8 T cells from ICI-A versus RA or PsA. **b,c)** Frequency of perforin^+^ cells, granzyme B^+^ cells (b) and Ki67^+^ cells (c) in CD8 T cells from RA or PsA SFMC cultured with IFN-*/3* or IFN-*y* for 16 hours or 72 hours. **d,e)** Representative flow cytometric plots showing expression of perforin and granzyme B (d) and Ki67 (e) in CD38^hi^ CD127^−^ (red) and CD38^−^ CD127^+^ (blue) CD8 T cell populations. * p<0.05 by Wilcoxon matched-pair test in (b,c).

